# Deep learning reveals a microprotein atlas in maize and uncovers novel regulators of seed amino acid metabolism

**DOI:** 10.1101/2025.11.14.688563

**Authors:** Anqiang Jia, Yawen Yang, Min Jin, Jimin Zhan, Mi Zhang, Sixuan Xu, Zhen Li, Keyu Tao, Yanhui Yu, Liying Feng, Jingxian Fu, Wenlong Li, Peng Zhang, Yongming Liu, Jieting Xu, Shouchuang Wang, Zuxin Zhang, Haiyang Wang, Jianbing Yan, Hai-Jun Liu

**Affiliations:** Yazhouwan National Laboratory, Sanya 572024, China; National Key Laboratory of Crop Genetic Improvement, Huazhong Agricultural University, Wuhan 430070, China; Hubei Hongshan Laboratory, Wuhan 430070, China; WIMI Biotechnology Co., Ltd., Qingdao 266100, China

## Abstract

Microproteins represent a vast and functionally important class of genes that remain largely unexplored in plant genomes. Here, we developed DeepMp, a deep learning framework that integrated multi-omics evidence and built the most comprehensive plant microprotein atlas to date, identifying 18,338 high-confidence candidates in maize. The majority of these appear to have originated *de novo* from regions previously annotated as noncoding, and they show hallmarks of rapid, lineage-specific evolution and pronounced tissue specificity. These novel microproteins were found integrated into core regulatory networks, particularly in organ development and nutrient storage. Focusing on the maize kernel, population-scale analyses linked microprotein expression to natural variation in amino acid content. We functionally validated three grain-filling-specific candidates originating from noncoding regions by CRISPR-Cas9 knockouts, which confirmed their roles as precise modulators of arginine, aspartate, and methionine levels, without pleiotropic effects on kernel morphology. Our study provides a foundational resource and analytical framework, establishes microproteins as a new and functionally important coding layer in maize, and uncovers a previously untapped source of targets for crop improvement.

## INTRODUCTION

The canonical view of a genome, with its neatly defined protein-coding genes, is incomplete. A growing evidence across all kingdoms of life has revealed that many of these hidden proteome encoded by small open reading frames (sORFs) that produce microproteins (∼5–100 amino acids)^1–4^. These are not just biochemical curiosities; they are functional players in metabolism^5^, development^6,7^, and stress responses^8,9^. However, their small size and frequent lack of sequence conservation have caused them to be systematically excluded from standard genome annotations, leaving a significant fraction of the functional coding landscape uncharted^10–12^.

In plants, this problem is particularly acute, however, comprehensive discovery remains challenging. Microproteins are typically short and expressed at low levels^13^, and plant genomes often feature the relative large sizes, repetitive nature^14,15^, and polyploid history of many crop genomes^16^ create a massive and challenging search space for sORFs, making conventional discovery methods inefficient. Current approaches which rely on peptidogenomics^17^ or homology^18^, both have intrinsic limitations: they either produce high false-positive rates or are blind to the novel, lineage-specific microproteins that appear to be a major feature of plant evolution^19,20^. A new strategy is clearly needed.

Deep learning models offer a powerful alternative^21,22^. For example, transformer-based models that leverage self-attention and large-scale unsupervised pre-training have increasingly been applied to peptide prediction tasks, including secreted-peptide identification^23,24^. Unlike homology-based approaches, they can learn the complex sequence features and contextual patterns of genuine coding sequences from multi-omics data, allowing them to predict novel genes with far greater accuracy^25^. Consequently, integrating multi-omics evidence with deep-learning methods represents a promising strategy for systematic microprotein discovery^21,22^, and this approach is more suitable to untangle the complexity of plant genomes.

We chose maize (*Zea mays*) as our test model. Its large (∼2.3 Gb), repeat-rich (more than 85%) genome makes it a formidable challenge for gene discovery^14,26^, while its immense genetic diversity and importance as a global crop make it an ideal system for translating genomic findings into functional understanding and agricultural application^27,28^. Furthermore, the systematic atlas of maize microproteins will establish a critical reference and paves the way for systematic characterization of these “hidden” genes in other major crops.

In this study, we present DeepMp, a deep-learning model to construct a comprehensive atlas of maize microproteins together with Ribo-seq and proteomic data from 704 samples. We characterized their evolutionary origins and expression patterns, then integrated this information with population-scale data to kernel development and amino acid metabolism. Furthermore, functional validation via CRISPR-Cas9 confirms three newly discovered microproteins, born from “non-coding” DNA, are precise regulators of seed amino acid composition. Our work provides both a powerful new framework for gene discovery and valuable novel targets for crop improvement.

## RESULTS

### A robust deep learning pipeline for genome-wide microprotein identification

To systematically identify microproteins (5–100 aa) in maize genome, we established an integrative framework that combines deep learning-based prediction with multi-omics validation (Figure 1A). The core of this pipeline is DeepMp, a hybrid deep learning model we designed to distinguish real microprotein-coding sequences from the vast background of non-coding sORFs.

**Figure 1.**
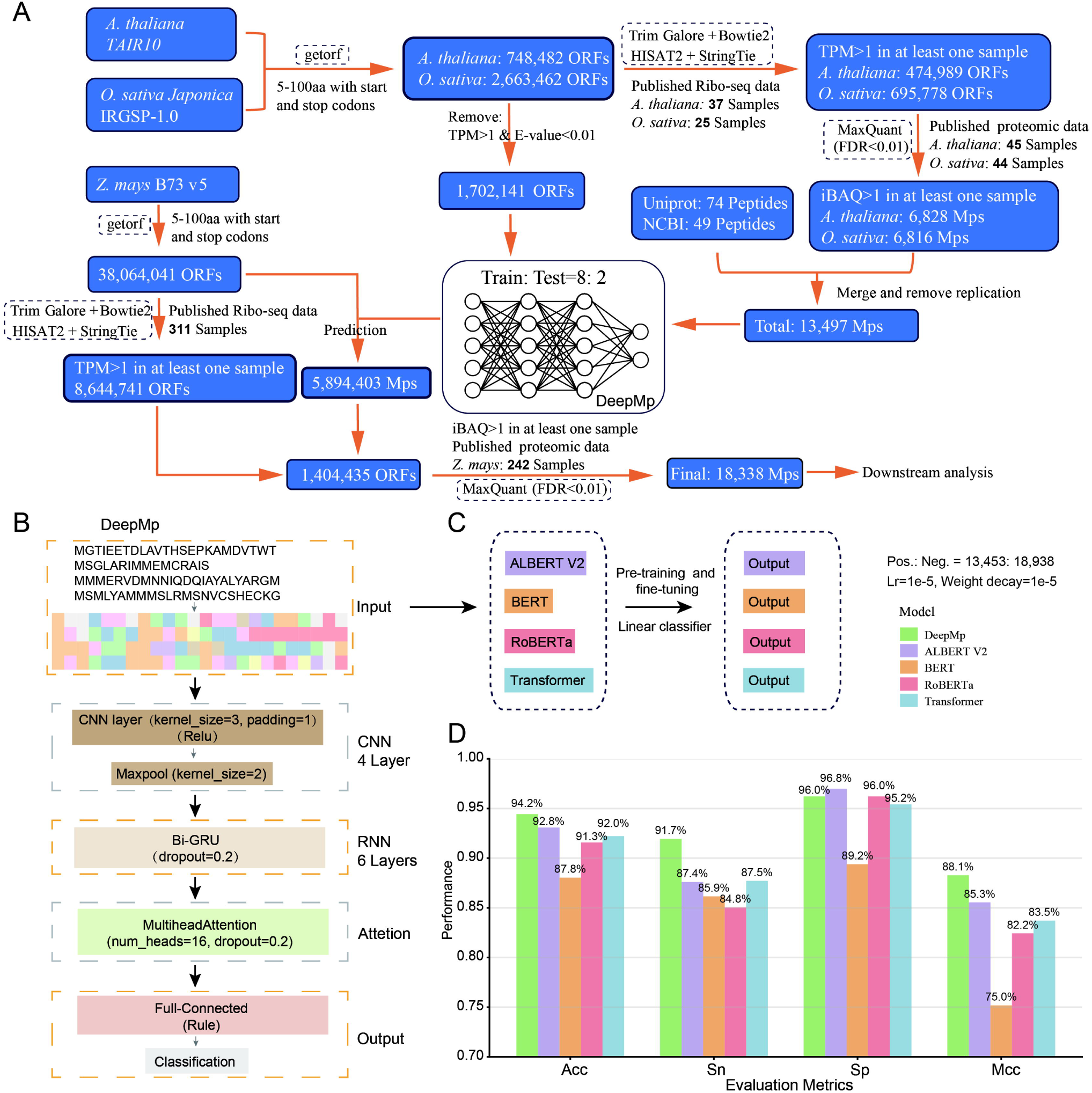
Computational pipeline and performance of DeepMp for microprotein identification. **(A)** Overview of the multi-omics and deep learning-based pipeline to identify high-confidence microproteins in maize. Software are enclosed in dashed boxes, and the sample cohort sizes for omics data are emphasized in bold. **(B)** Architecture of the DeepMp model, a hybrid network integrating convolutional neural networks (CNN), bidirectional gated recurrent units (Bi-GRU), and multi-head self-attention mechanisms. Input sequences were encoded (amino acids 1–20) and zero-padded to a length of 100 residues. **(C)** Benchmarking of DeepMp against four pre-trained language models. All models were fine-tuned on an identical balanced dataset (13,453 positive and 18,938 negative sequences) under consistent hyperparameters (Lr, learning rate: 1e-5; Weight decay: 1e-5) to ensure a fair comparison. **(D)** Performance comparison on the held-out test set. DeepMp demonstrates superior performance. Metrics include: Acc, Accuracy; Sn, Sensitivity; Sp, Specificity; MCC, Matthews correlation coefficient.

To train DeepMp, we first built a high-confidence reference set of 13,497 non-redundant microproteins (5–100 aa), which was built upon 6,828 and 6,816 entries from *Arabidopsis* and rice that were rigorously validated by extensive Ribo-seq (37 *Arabidopsis* and 25 rice) and proteomic data (45 *Arabidopsis* and 44 rice) (Figure 1A; Tables S1–S2), and subsequently expanded with 123 peptides from public databases. The model, which combines convolutional and recurrent neural network layers with a multi-head attention mechanism, was trained on the intrinsic sequence features of these validated microproteins (Figure 1B; see STAR Methods).

In head-to-head comparisons, DeepMp significantly outperformed established language models (ALBERT V2, BERT, RoBERTa, and Transformer), achieving a superior balance of accuracy (Acc = 0.94) and sensitivity (Sn = 0.92), with a highest Matthews correlation coefficient (measures how well the predictions match the actual labels, MCC = 0.88) (Figures 1C–1D). This high performance held even on imbalanced datasets (stable Acc and AUC for positive:negative ratio ≈ 1:100; Figures S1A–S1B). We also observed high similarity in features between the positive set and the predicted microproteins, such as residue composition, hydrophobicity, and amphipathicity, as well as distinct characteristics from the negative set (Figure S1C). These results collectively underscored our model’s predictive power and confirmed its robustness for genome-scale prediction.

We next applied the framework to the maize reference genome (B73_v5). From an initial pool of over 38 million possible sORFs, DeepMp predicted nearly 5.9 million candidate microproteins. We then subjected these predictions to stringent experimental filters, retaining only those with direct evidence of translation from 311 Ribo-seq experiments (transcripts per million, TPM > 1; Table S1) and peptide-level confirmation (intensity-based absolute quantification, iBAQ > 1; FDR < 0.01; Table S2) from 242 proteomic datasets. This rigorous, multi-tiered approach yielded 18,338 high-confidence maize microproteins (Figure 1A), representing the most comprehensive atlas of this gene class in any plant to date.

### The maize microprotein atlas reveals a shadow proteome of *de novo* origin

The 18,338 identified microproteins constitute a novel and diverse class of genes. They are predominantly small, with a peak length distribution between 45–75 amino acids (Figure 2A). Strikingly, the majority (76.1%) lacked any recognizable protein domain, and most of them (98.4%) lacked a secretion signal (Figure 2B), suggesting they likely function intracellularly. They can be clustered (50% identity and 80% coverage thresholds) into 11,605 distinct families (Figure S4A), indicating a rich reservoir of non-canonical peptides with potentially functional diversity.

**Figure 2.**
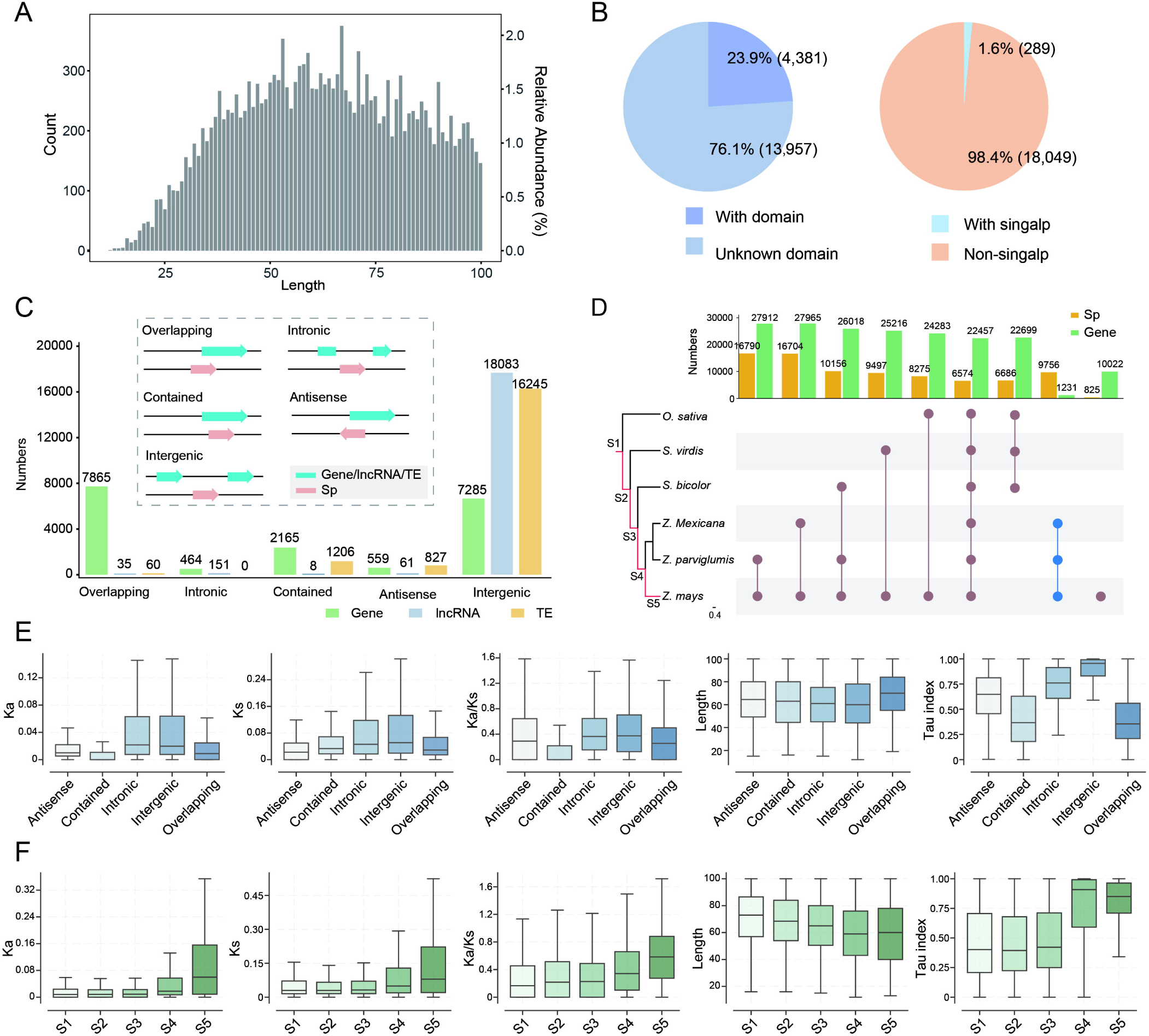
Genomic and evolutionary characteristics of maize microproteins. **(A)** Length distribution of amino acid sequence for identified microproteins. **(B)** Proportions of microproteins containing a recognizable Pfam domain or a predicted signal peptide. **(C)** Genomic origins of microproteins relative to annotated protein-coding genes, long non-coding RNAs (lncRNAs), and transposable elements (TEs). **(D)** Conservation of maize microproteins and protein-coding genes across five Poaceae species, assessed by sequence homology and genomic synteny. The Upset plot illustrates shared and species-specific microprotein families, with conservation strata (S1–S5) defined on the left. Blue connecting line highlights families conserved only between maize and its teosinte progenitors. (**E–F**) Boxplots comparing evolutionary features of microproteins across different genomic contexts (F) and conservation strata (G, corresponding to panel E). The analyzed metrics include the rates of nonsynonymous (Ka) and synonymous (Ks) substitution, the Ka/Ks ratio, protein length, and the tissue-specificity index (Tau).

Genomically, these microproteins are not randomly distributed. They are enriched in the gene-dense chromosomal arms and depleted near centromeres (Figure S2A). Critically, a large proportion (42.9%) overlaps with annotated protein-coding genes, followed by those from intergenic regions (39.7%) (Figure 2C). A minority were fully embedded within genes (11.8%), while smaller subsets were derived from antisense strands (3.0%) or resided entirely within introns (2.5%). Only limited were overlapped with the current annotated long non-coding RNAs (lncRNAs, 0.2%) and transposable elements (TEs, 0.3%), respectively. This reminded us that, even within the space of annotated protein-coding genes, coding diversity may have been significantly underestimated. The localization within previously designated non-coding regions, combined with their lack of canonical domains, strongly suggests that many of these microproteins are products of *de novo* gene birth. Further evolutionary analysis supported the *de novo* birth hypothesis. While a core set (35.8%) of microproteins showed conservation predating the 49 million years of grass divergence (Figure 2D), over half (53.2%) were *Zea*-genus specific, with 4.5% exclusively found in *Zea mays*. This represented an eightfold enrichment of lineage-specificity compared to canonical protein-coding genes, whereas only 2.4% were unique to *Zea* genus. This recent and explosive expansion pointed to microproteins as a major source of evolutionary innovation in the *Zea* lineage.

### Younger microproteins evolve rapidly and exhibit tissue-specific expression

We next investigated the evolutionary dynamics of these newly identified microproteins by integrating selection pressure (Ka/Ks) with transcriptomic profiles across 56 tissues and developmental stages (Table S3). Microproteins arising from non-coding (intronic and intergenic) regions exhibited markedly relaxed purifying selection, as evidenced by their significantly higher Ka/Ks ratios compared to those overlapping existing genes (Figure 2E). This relaxed selection is coupled with a striking pattern of tissue-specific expression, with intergenic microproteins showing the highest degree of tissue specificity (Figures 2E and S3A). This pattern is a typical signature of neofunctionalization, where new genes are initially expressed in narrow developmental or environmental niches before potential integration into broader regulatory networks^29,30^.

This trend is mirrored when we stratify microproteins by evolutionary age. Phylogenetic analysis classified microproteins into five evolutionary strata (Figure 2D), 3,113 microproteins originated prior to the *O. sativa*-*Zea* ancestral divergence (S1), 4,417 emerged before the *S. viridis*-*Zea* divergence (S2), 3,300 predated the *S. bicolor*-*Zea* divergence (S3), 6,683 arose before *Zea*-specific divergence (S4), and 825 specific to *Z. mays* (S5) (Figure S2C). Younger microproteins, particularly those that arose within the *Zea* genus (strata S4–S5), were evolving more rapidly (higher Ka/Ks) and shorter, and exhibit greater tissue specificity than more ancient ones (Figures 2F and S3B). Despite their rapid sequence evolution, predicted 3D structures show that they maintained plausible protein-like folds (mean pLDDT > 60%; pLDDT: predicted Local Distance Difference Test), rich in alpha-helices and coils (Figure S4). Together, these data painted a picture of a highly dynamic gene class, where *de novo* birth from non-coding DNA provided a reservoir of rapidly evolving, tissue-specific regulators^31–33^.

### Co-expression networks revealed microproteins in kernel development

To infer the biological functions of these uncharacterized microproteins, we constructed a global co-expression network using transcriptomic data from 56 tissues, comprising 26,950 protein-coding genes and 11,429 microproteins. Microproteins were not isolated but integrated into 25 distinct functional modules alongside known protein-coding genes (Figures 3A and S5–S6). Seven modules exhibited significant correlations (Pearson’s r ≥ 0.6, p < 0.01) with seed, leaf, or root tissues, suggesting tissue-specific functional specialization (Figures 3A and S6). Among these, four modules were strongly associated with seed development, implicating microproteins in processes from carbohydrate and lipid storage to organ morphogenesis (Figure 3A). Other modules linked microproteins to photosynthesis in leaves and cell wall biosynthesis in roots, demonstrating their system-wide integration (Figure S6).

**Figure 3.**
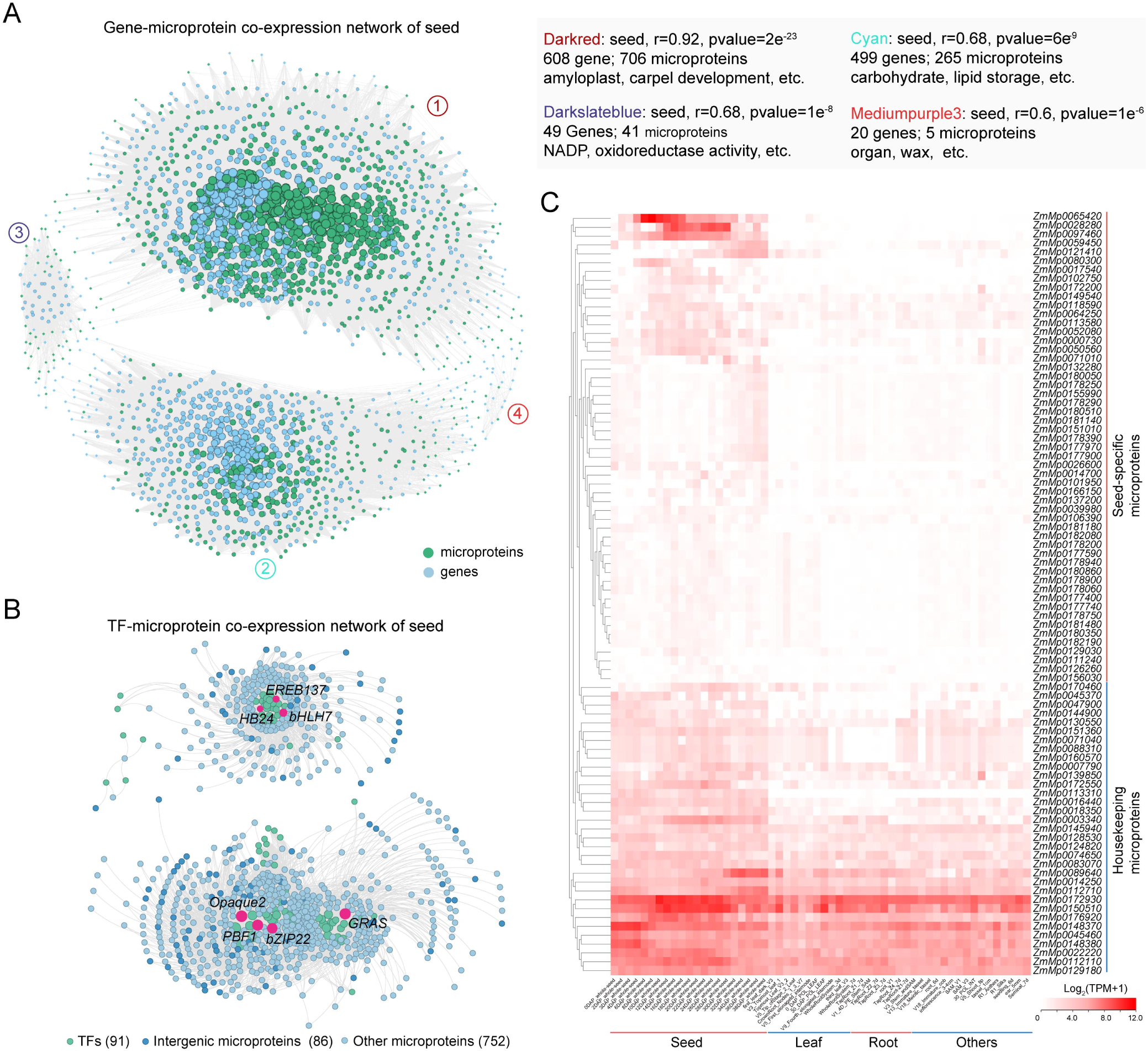
Co-expression network of microproteins and genes. **(A)** Four co-expression modules significantly associated with the Seed. The Pearson correlation coefficient (r) and associated p-values are indicated at the top right of the figure, along with the top enriched Gene Ontology (GO) terms. Genes are represented by light blue circles, and microprotein by green circles. The numbered circles adjacent to each module (left) correspond to the colored module names at the top right. **(B)** Subnetwork of the seed-specific module, depicting regulatory interactions between hub transcription factors (TFs, green circles) and microproteins (blue circles). The intergenic microproteins are colored in dark blue and the others in light blue. The network comprises 91 TFs, 86 intergenic microproteins, and 752 microproteins from other genomic regions. The size of each circle represents the number of co-expressed genes. **(C)** Expression heatmap of the 86 intergenic microproteins across 56 diverse maize tissues and developmental stages. Expression levels are displayed as log_2_(TPM+1).

To pinpoint their specific roles in seed development, we constructed a more focused network centered on transcription factors (TFs). TF-microprotein co-expression network revealed that hundreds of microproteins, comprising 86 intergenic and 752 non-intergenic microproteins, were co-expressed with 91 key regulators of kernel development and nutritional quality (Figure 3B). This included *Opaque2*, *PBF1*, *GRAS11*, and *bZIP22*, which are known to coordinately control zein storage protein expression^34,35,36,37^, and several other TFs (*bHLH7*, *EREB137*, *HB24*, *ZFP2*, and *NF-YB7*) specifically expressed during seed developmental stages^38^ (Figure S7). Remarkably, many of these network-associated microproteins, particularly those located in intergenic regions, demonstrated expression patterns that peaked specifically during grain-filling stages (Figure 3C). Together, these results demonstrate that microproteins are incorporated into the core transcriptional circuitry likely participating in maize kernel development and protein quality regulation.

### Genetic association of microproteins with kernel amino acid content

The network analyses suggested a role of microproteins in kernel quality. To test this genetically, we correlated microprotein expression in another 354 maize inbred lines at 15 days after pollination (DAP) with the contents of 18 amino acids, and kernel-related yield traits. This identified 575 intergenic microproteins whose expression was significantly associated with at least one trait (|r| ≥ 0.15, p < 0.05). Among these, 22 microproteins that overlap the TF-microprotein network showed significant correlations with 14 amino acids, including aspartic acid (Asp), lysine (Lys) and arginine (Arg), as well as with kernel dimensions (length and thickness) (Figures 4A–4B). Subsequent eQTL mapping revealed 10 of these microproteins were cis-regulated (FDR < 0.05; peak SNPs located within ±2 kb of the transcriptional unit) (Figures 4C and S8). Multi-tissue expression profiling further prioritized three microproteins (*ZmMp0003340*, *ZmMp0080300*, and *ZmMp0172550*) with specific expression during grain-filling (Figures 3C and S9). Genomic annotation placed *ZmMp0003340* within the 3′ UTR of *Zm00001eb009010*, *ZmMp0080300* in an intergenic locus, and *ZmMp0172550* in the 5′ UTR of *Zm00001eb421180*. Each of these three microproteins was excluded as TE/lncRNA/miRNA-derived and was supported by both transcriptomic and translatomic evidence (Figure 4D). Haplotype analysis based on the *cis*-eSNPs revealed a crucial pattern: different alleles of these microprotein genes were associated with significant differences in the levels of specific amino acids, but had no significant effect on kernel size or weight (Figures 4E–4H and S10). This genetic evidence strongly suggests these three microproteins likely act as specific modulators of amino acid metabolism.

**Figure 4.**
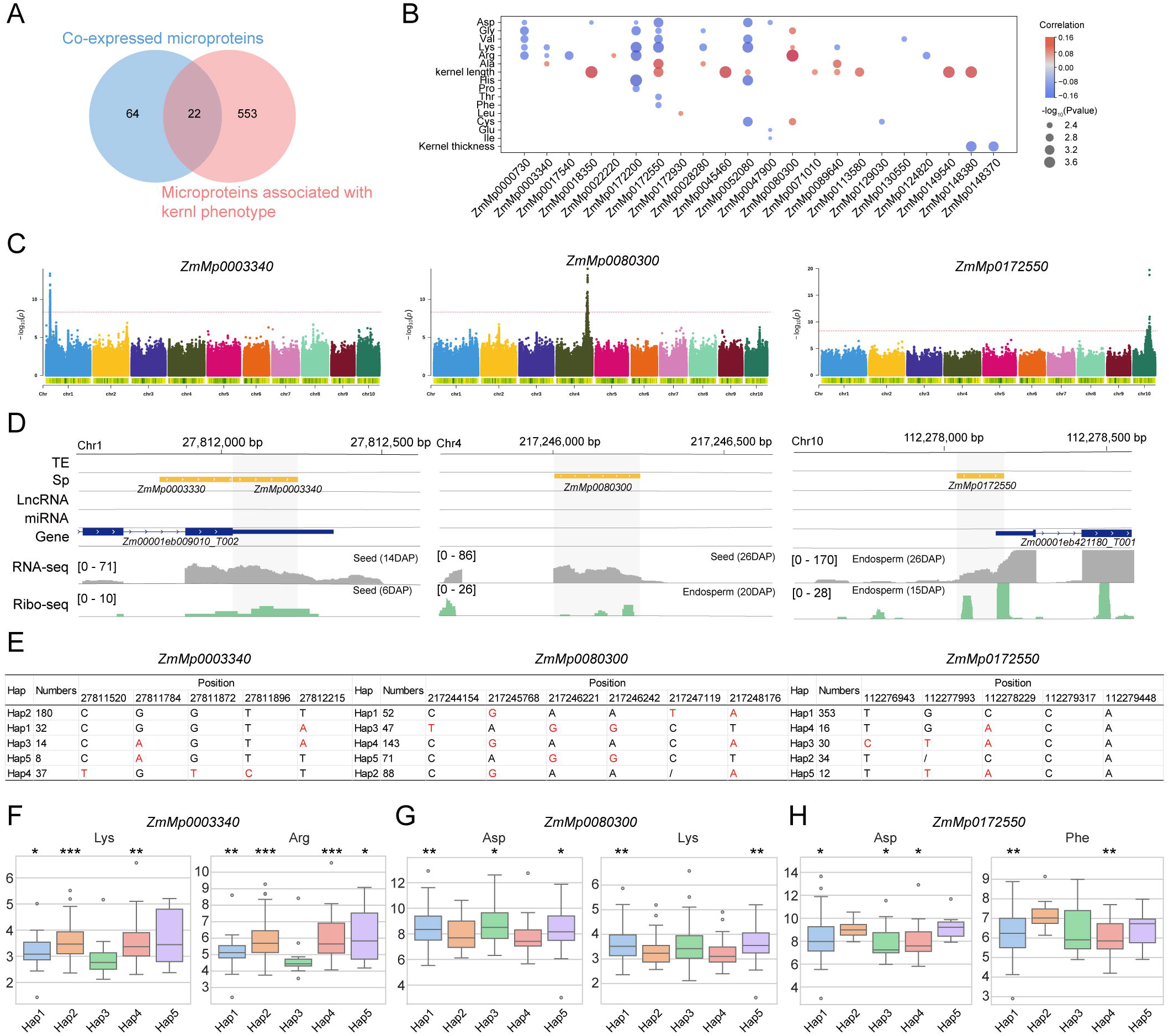
eQTL mapping and haplotype analysis identify three microproteins associated with seed amino acid content. **(A)** Venn diagram illustrating the strategy to prioritize candidate microproteins, selecting those present in seed-associated co-expression modules, linked to an eQTL, and correlated with kernel phenotypes. **(B)** Bubble plot showing the Pearson correlation between the expression of 22 candidate microproteins and kernel amino acid content and kernel size traits. Red and blue indicate positive and negative correlations, respectively. Bubble size corresponds to statistical significance (−log_OO_(P-value)). **(C)** Manhattan plots of eQTL analyses for the expression of three microproteins. The genome-wide significance threshold (red dashed line) was defined by a significance threshold after 5% Bonferroni correction. **(D)** Genome browser view showing transcriptional (RNA-seq) and translational (Ribo-seq) evidence for the three microprotein loci, along with their genomic context relative to neighboring genes, lncRNAs, TEs, and miRNAs. **(E)** Haplotype blocks formed by significant eQTL-associated SNPs within 2-kb-flanking regions of each microprotein. **(F–H)** Association between haplotypes for each of the three microproteins and the relative content of specific amino acids in mature seeds. Data are presented as mean ± SD. Statistical significance was determined by the Kruskal-Wallis test (*P < 0.05, **P < 0.01, ***P < 0.001).

### Evolution and translational activity of the three candidate microproteins

The three prioritized candidates represent compelling evolutionary case studies. ZmMp0003340 and ZmMp0080300 appear to be *de novo* originated following the divergence of the *S. bicolor* and *Zea* lineages, whereas ZmMp0172550 was specific to the *Zea* lineage (Figure 5A). While ZmMp0003340 maintains strict single-copy conservation, although *S. bicolor* orthologs carry an N-terminal variant with a C-terminal extension relative to *Zea* counterparts (Figures 5A–5B), ZmMp0080300 arose from a gene duplication event in *Zea*. The ancestral copy retained a Defensin domain (Embryo Surrounding Region 6, ESR6, a known embryo-surrounding region protein involved in kernel development^39^) and a signal peptide, while the new copy lost the domain and evolved into a microprotein, a clear case of neofunctionalization (Figures 5A–5B). Comparative genomic analysis revealed polymorphic loss of the signal peptide in ZmMp0080300 across modern maize population, indicating ongoing functional diversification (Figure S11). The promoters (2 kb upstream regions) of all three contain cis-regulatory elements associated with kernel development (O2-site, RY-elements, GCN4_motif) and hormone response (GARE-motif, P-box, TCA-element, TGA-element etc.) (Figures 3B, 4F–4H and 5C), consistent with their expression patterns and potential functional roles.

**Figure 5.**
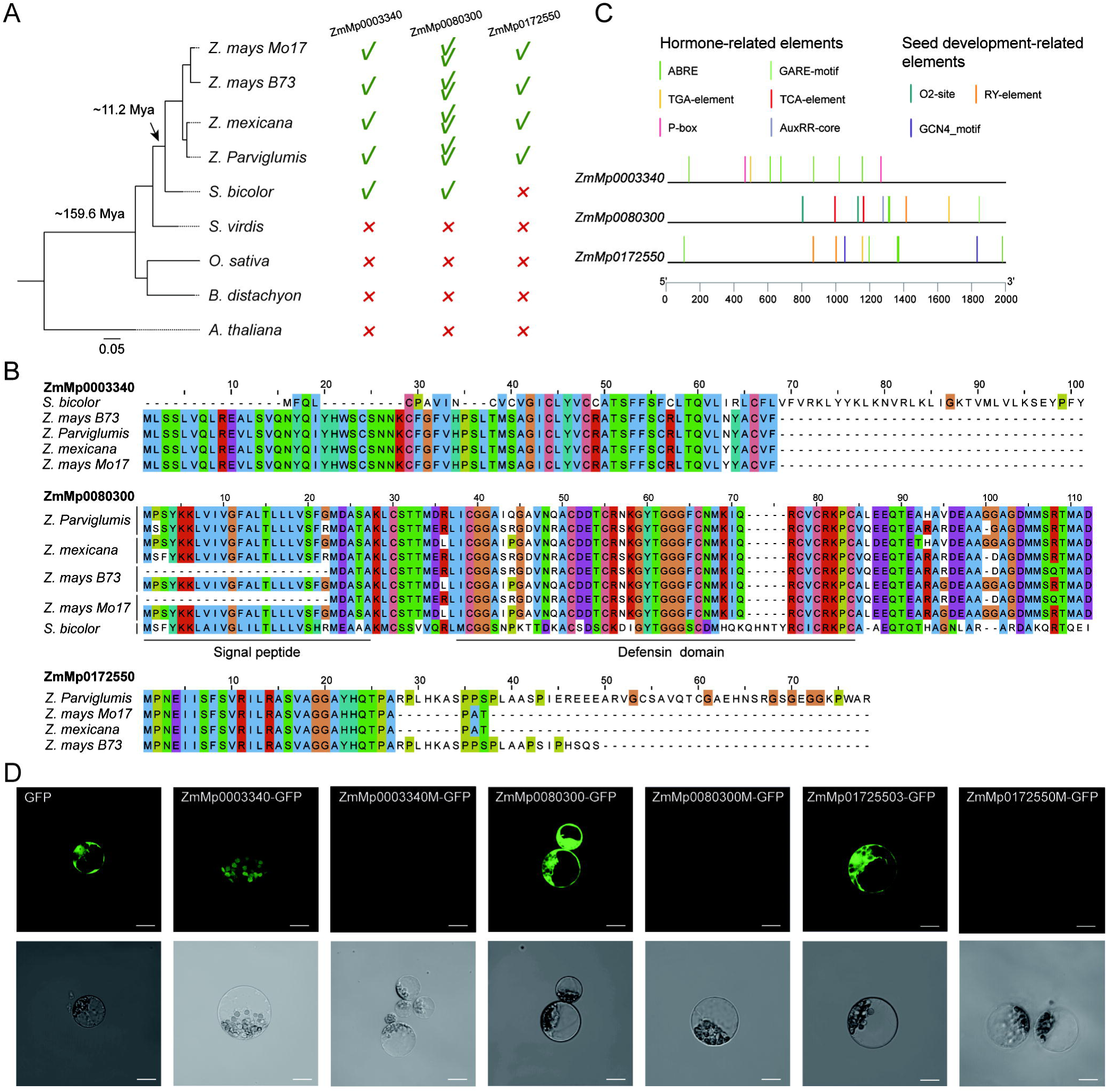
Evolutionary and functional characterization of the three microproteins. **(A)** Distribution of homologs for the three microproteins across eight plant species. Presence (√), absence (×) and copy number (multiple √) are indicated. **(B)** Multiple sequence alignment of amino acid sequences for the three microproteins and their orthologs, highlighting conserved residues. **(C)** Promoter analysis (2 kb upstream) reveals an enrichment of cis-regulatory elements associated with hormone signaling and seed development. **(D)** Validation of microprotein translation in maize protoplasts. Each microprotein was fused to a C-terminal GFP reporter. Robust GFP fluorescence confirms in vivo translation, with free GFP serving as a positive control. Scale bar, 10 µm.

To validate their translational activities, we employed a dual-luciferase reporter assay in maize protoplasts based on our previous protocol^7^: experimental constructs carried mutated start codons in both the microprotein (ATG→ATT) and GFP (ATG→CTT), while control constructs contained mutations only in GFP. Constructs containing the native microprotein sORFs drove robust reporter expression, while mutating the start codon (ATG→ATT) completely abolished translation (Figure 5D). These results confirmed that these sequences are not only transcribed but are actively translated into peptides *in planta*.

### CRISPR-Cas9 confirms microprotein roles as fine-tuners of seed amino acid metabolism

To definitively assess the *in vivo* function, we generated homozygous CRISPR-Cas9 knockout mutants for all three candidate microproteins in the maize inbred line KN5585, each with two independent events to ensure phenotypic reproducibility (Figure 6A). Given their kernel-specific expression (Figure S9), we first examined kernel morphology. Remarkably, null mutations in any of the three microproteins resulted in no discernible phenotype in kernel size or overall morphology (Figures 6B–6C). Kernels in all mutant lines were in distinguishable from wild-type.

**Figure 6.**
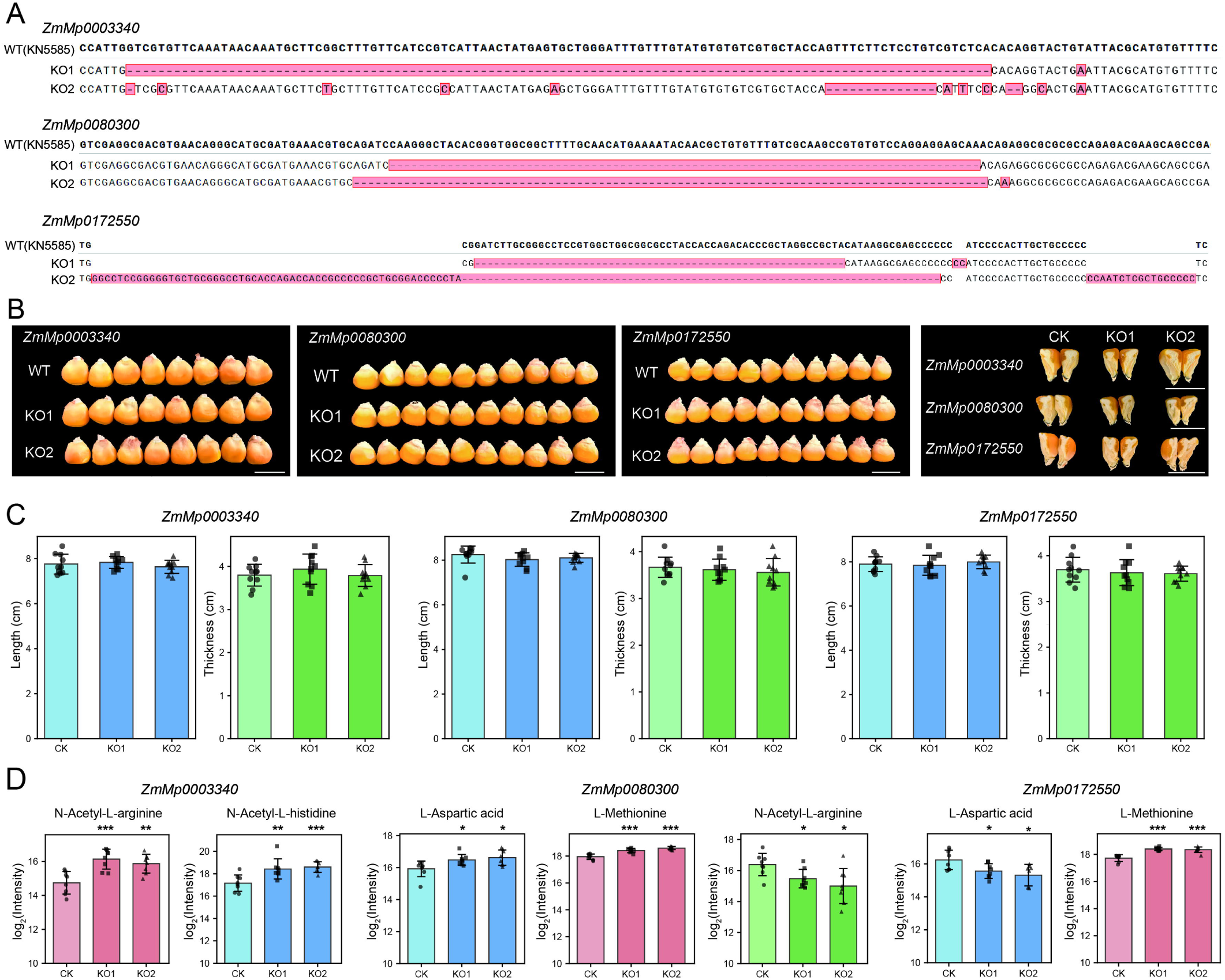
CRISPR-Cas9-mediated knockout of microproteins alters seed amino acid composition. **(A)** Schematic of the CRISPR-Cas9 target sites and induced mutation alleles. Two independent homozygous mutant alleles are shown for each gene, with red boxes indicating the extent of genomic deletions. **(B)** Representative images of mature kernels from wild-type (WT) and homozygous mutant plants, showing no discernible difference in external morphology or internal structure (longitudinal sections). **(C)** Quantification of kernel length and thickness for the mutants corresponding to the three microproteins. Data are presented as mean ± SD (n=10). **(D)** Relative content of specific amino acids in mature kernels of WT and mutant lines. Data are mean ± SD (n=8). Significant differences from WT were determined by a two-tailed Student’s t-test (*P < 0.05, **P < 0.01, ***P < 0.001).

Liquid chromatography mass spectrometry (LC-MS) analysis of mature kernels revealed their roles in amino acid metabolism. Consistent with the natural variations identified earlier (Figures 4F–4H), knockout of *ZmMp0003340* led to a marked increase of N-Acetyl-L-arginine, implicating its involvement in arginine metabolism (Figures 6D and S12A). Similarly, the *ZmMp0080300* and *ZmMp0172550* mutants elicited opposing effects on L-Aspartic acid levels, with the former significantly increasing and the latter decreasing them, respectively (Figure 6D). Intriguingly, our analysis uncovered additional regulatory roles beyond these predicted pathways: *ZmMp0003340* knockout increased N-Acetyl-L-histidine abundance, *ZmMp0080300* knockout elevated L-Methionine and N-Acetyl-L-arginine levels, and the knockout of *ZmMp0172550* affected the levels of L-Methionine and total protein content in seeds (Figures 6D and S12B), suggesting broader regulatory functions and potential functional redundancy or crosstalk in amino acid metabolism^40,41^.

Collectively, our functional dissection uncovers a striking phenotypic dichotomy: these *de novo*, kernel-specific microproteins are not required for organogenesis or biomass accumulation, but instead, they act as precise modulators of nutritionally critical amino acids, like arginine, aspartate^42^, and methionine^43^ in the seeds.

## DISCUSSION

The systematic identification of microproteins in plants, particularly in species with large and complex genomes like maize, has long been a formidable challenge. The sheer size of the potential search space, over 38 million sORFs in the maize genome, creates a significant computational and statistical barrier, leading to high false-positive rates and obscuring the discovery of genuine, functional peptides. Current efforts have primarily focused on identifying microproteins encoded by lncRNAs^44^, characterizing isolated functional mechanisms^45,46^, and exploring small and other noncanonical peptides^47–50^. However, systematically uncovering microproteins in complex plant genomes remains a formidable challenge. To address these limitations, we developed an integrated pipeline that represents a paradigm shift for microprotein discovery in complex genomes.

By combining our deep learning model, DeepMp, in evolutionarily conserved features and coupling its predictions with multi-omics validation at an unprecedented scale, including transcriptomics, translatomics (Ribo-seq), and proteomics, we established a robust framework for high-confidence identification (Figure 1A). Our strategy yielded an atlas of 18,338 high-confidence microproteins in maize, representing the most comprehensive catalog in any plant to date. This work not only substantially expanded the annotated coding capacity of maize, but also provides a foundational resource for the entire community. Notably, the 1.4 million translated sORFs (Figure 1A) identified by Ribo-seq but not yet confirmed by proteomic represent a rich reservoir for future functional mining, as proteomic depth continues to increase.

Our evolutionary analyses revealed that microproteins are a major driver of genetic innovation in maize. An impressive 53.2% of the identified microproteins were specific to the *Zea* genus, which is a significantly higher rate of *de novo* emergence than that of canonical protein-coding genes (∼2.4%). These young, often intergenic, microproteins exhibit classic signatures of neofunctionalization: under relaxed purifying selection (as evidenced by higher Ka/Ks ratios) and display highly tissue-specific expression patterns (Figures 2F–2G and S3). These findings support the hypothesis that non-coding DNA acts as a crucible for evolutionary experimentation, generating new peptides that can be rapidly integrated into existing biological networks^7^. Our co-expression analyses support this model, by demonstrating that microproteins were involved into the transcriptional circuitry that governs core agronomic traits (Figures 3A-3B), including kernel development where they were co-regulated with master regulators like *Opaque2* (Figure S13).

Importantly, our study bridges the gap from genome-wide discovery to validated biological function and agronomic relevance. By integrating co-expression networks, population-level expression data (eQTL) with phenotypic associations, we identified and validated three candidate microproteins in amino acid metabolism and had no discernible impact on kernel morphology or yield components (Figures 6B–6D). This discovery of a functional decoupling between metabolism and growth is highly significant for crop improvement. It suggests that microproteins can act as metabolic fine-tuners, offering a new class of targets for enhancing nutritional quality without the risk of yield penalties that often accompany mutations in pleiotropic developmental genes.

In summary, this study systematically charted the hidden coding landscape of the maize genome. We demonstrated that microproteins were not a minor curiosity but a large and dynamic class of genes that likely have played a pivotal role in the recent evolution of maize. By establishing their integration into key agronomic pathways and demonstrating their potential as precise targets for metabolic engineering, our study redefined the boundaries of the coding genome in crops. It provided not only a rich resource but also a clear strategy for harnessing this previously hidden layer of genetic diversity to accelerate crop genetic improvement.

### Limitations of the study

While our study established a foundational atlas and a robust discovery framework, its scope and methodology present clear limitations that define critical next steps for the field. First, DeepMp was trained on a reference set of conserved microproteins from limited taxa. Consequently, it is inherently biased against the most evolutionarily novel, lineage-specific candidates. Second, our validation relied on a large but finite set of Ribo-seq and proteomic experiments. While extensive, these datasets cannot capture the entire spatiotemporal expression landscape. Microproteins expressed only under specific environmental stresses or during narrow developmental windows may remain undetected in our current atlas. Third, our functional characterization was a proof-of-concept, focused on three candidates implicated in kernel amino acid metabolism. The molecular mechanisms for the vast majority of the over 18 thousands of microproteins in our atlas remain to be elucidated. This atlas should therefore be viewed as a rich resource for systematic, hypothesis-driven functional genomics. Finally, the functional inferences drawn from co-expression networks are, by nature, correlational. They provide powerful hypotheses about transcriptional modules but do not resolve the underlying molecular mechanisms. Dissecting whether these microproteins function via direct protein-protein interactions with network partners, enzymatic activity, or other post-translational modes will require further targeted biochemical approaches.

## RESOURCE AVAILABILITY

### Lead contact

For further information or requests for resources and reagents, please contact Anqiang Jia or Hai-Jun Liu.

### Materials availability

This study did not generate any unique reagents. All outputs from the analyses are listed in the “Data and Code Availability” section below. All data generated in this study, including expression results of microproteins and genes, are available on figshare (https://figshare.com/account/articles/30228853).

### Data and code availability

This study analyzed publicly available datasets, the accession numbers of which are provided in the Key Resources Table. Data generated during this study’s downstream analyses have been deposited in figshare and will be publicly available upon publication.

All original code has been deposited on GitHub and will be publicly accessible as of the date of publication (https://github.com/jiaanqiang/DeepMp). Any additional information required to reanalyze the data reported in this study is available from the lead contact upon request.

## ACKNOWLEDGMENTS

We are grateful to all members of the Maize Genome and Breeding Team at Yazhouwan National Laboratory for their insightful discussions and valuable feedback throughout this study. We also extend our thanks to Junjie Zhou and Jing Li from the Agricultural Bio-Multi-Omics Platform Team at the same Laboratory for their technical support in kernel metabolite analysis. This work was supported by the Hainan Postdoctoral Research Project (JB24BYKY02 to J.A.) and the Key Research Project of Guangdong Province (2022B0202060005 to H.W.). Computational resources were provided by the Advanced Computing Center of Yazhou Bay Science and Technology City.

## AUTHOR CONTRIBUTIONS

A.J., Y.Y., H.W., J.Y. and H.-J.L. designed the research, analyzed the data, drafted the manuscript, and prepared the figures and tables. Y.Y., J.X., and A.J. performed maize microprotein knockouts, screened positive individuals, and conducted phenotypic analysis. M.J., M.Z. and A.J. measured the amino acid content of kernels. A.J., J.Z., and Z.L. managed field planting and harvesting. S.X. and A.J. carried out maize protoplast transformation experiments. H.Y. provided guidance and methodologies for microprotein functional studies and activity validation. L.F., J.F., W.L., and P.Z. assisted with preparing seed samples and analyzing data. K.T. measured the seed protein content. Y.L., S.W., Z.Z., and J.Y. contributed to revising the manuscript. H.W., J.Y. and H.-J.L. supervised the research, coordinated data production, and finalized the manuscript with input from all authors. All authors have read and approved the final version.

## STAR□METHODS

### KEY RESOURCES TABLE

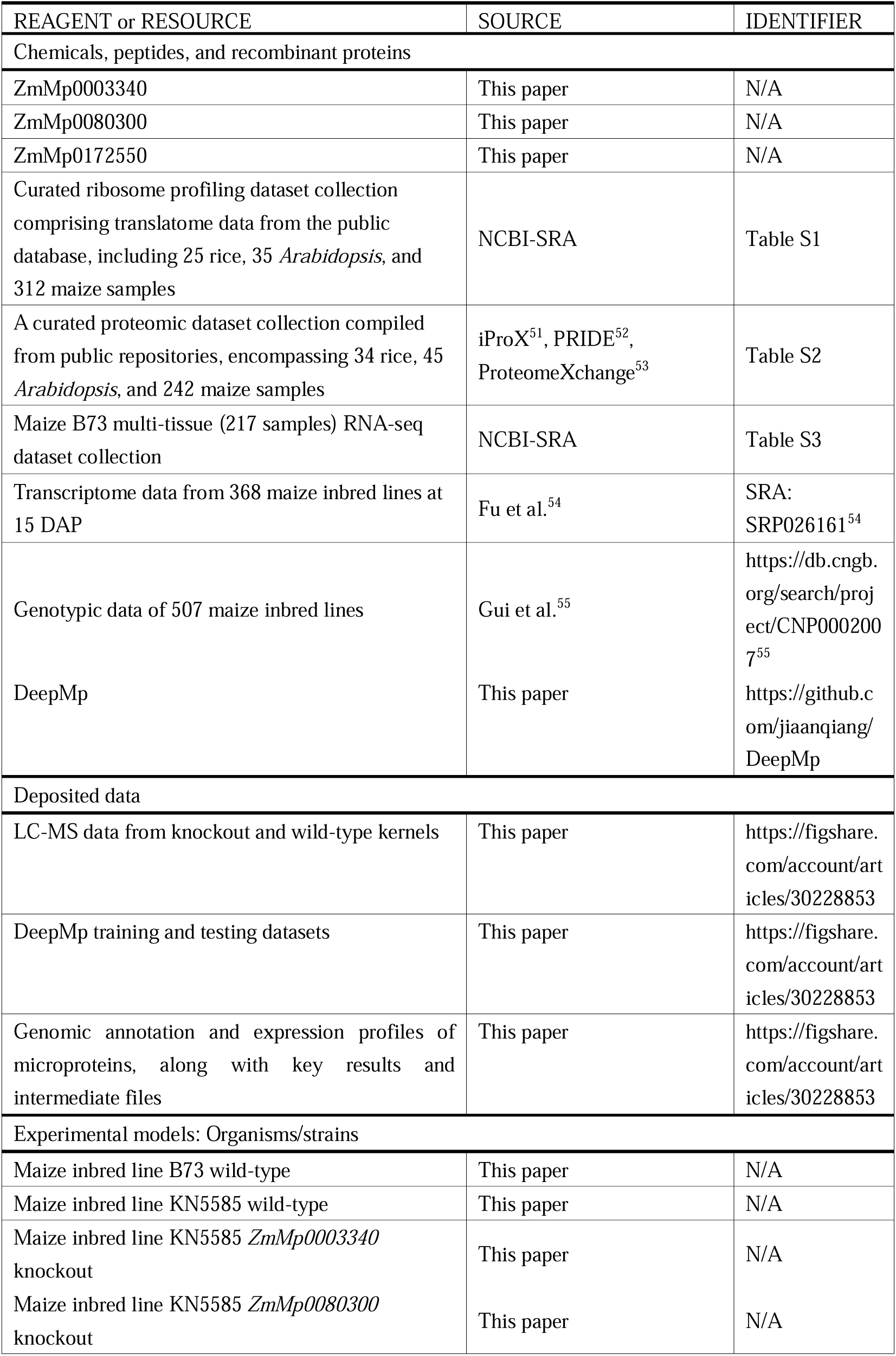
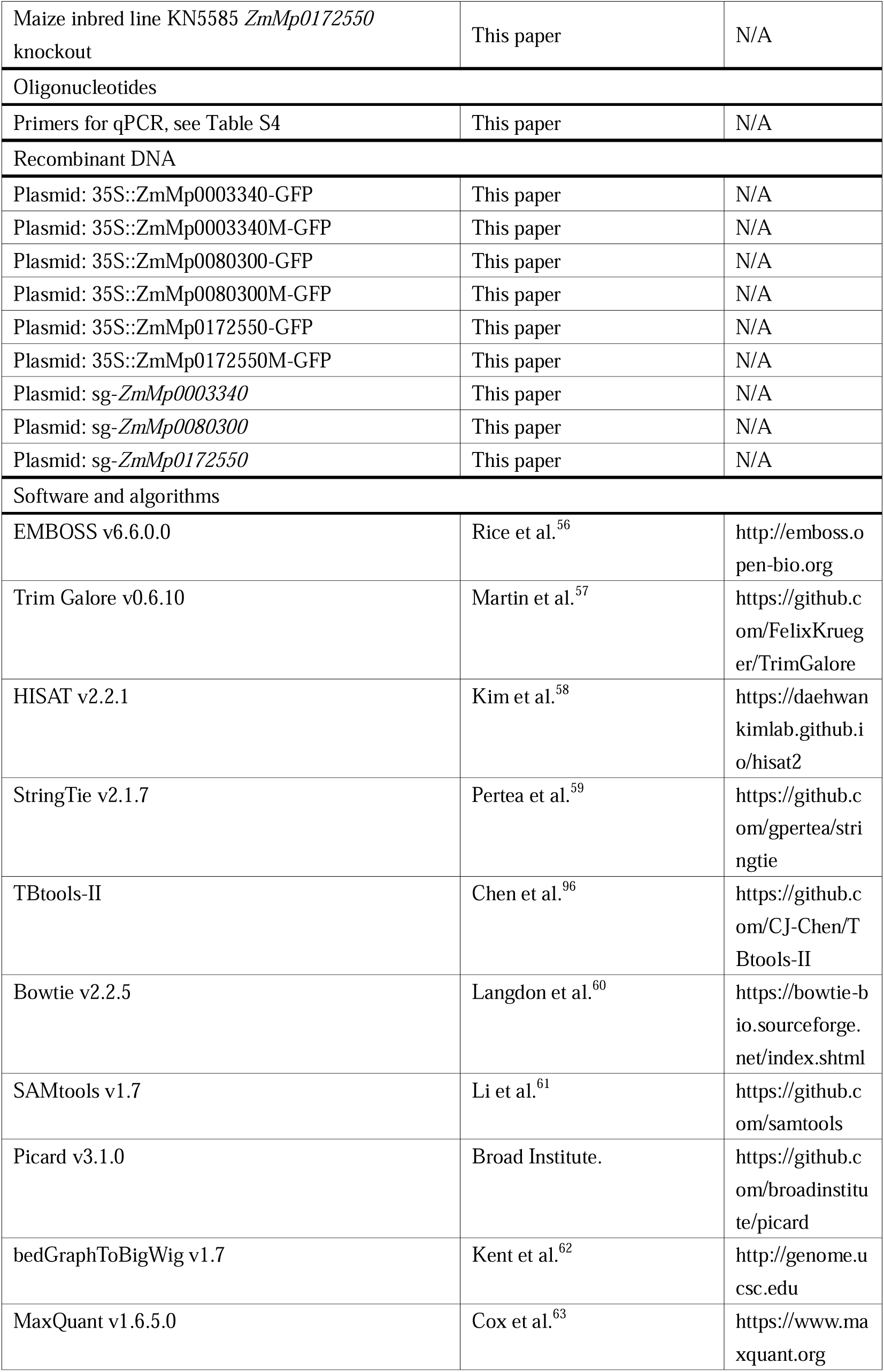
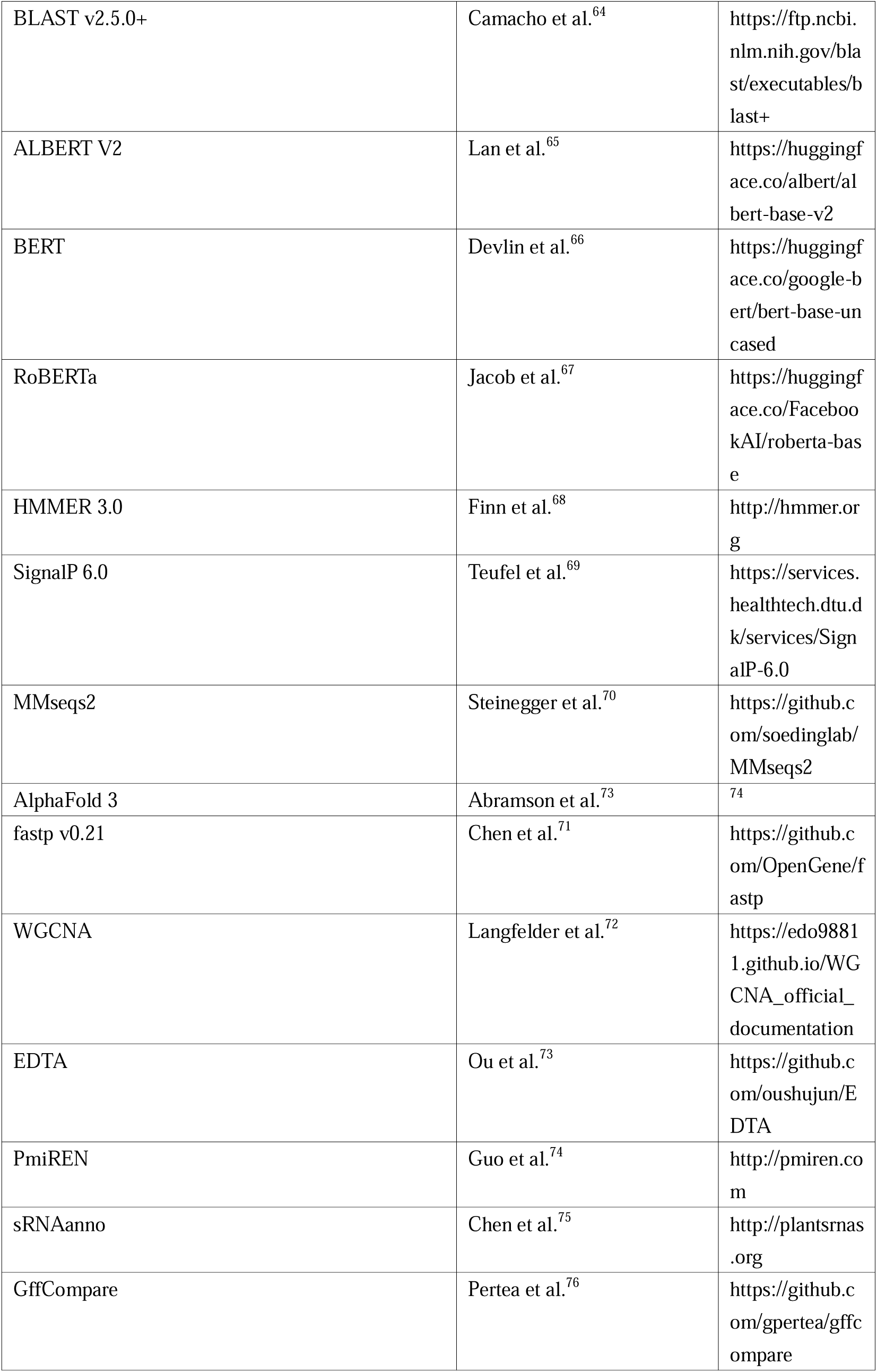
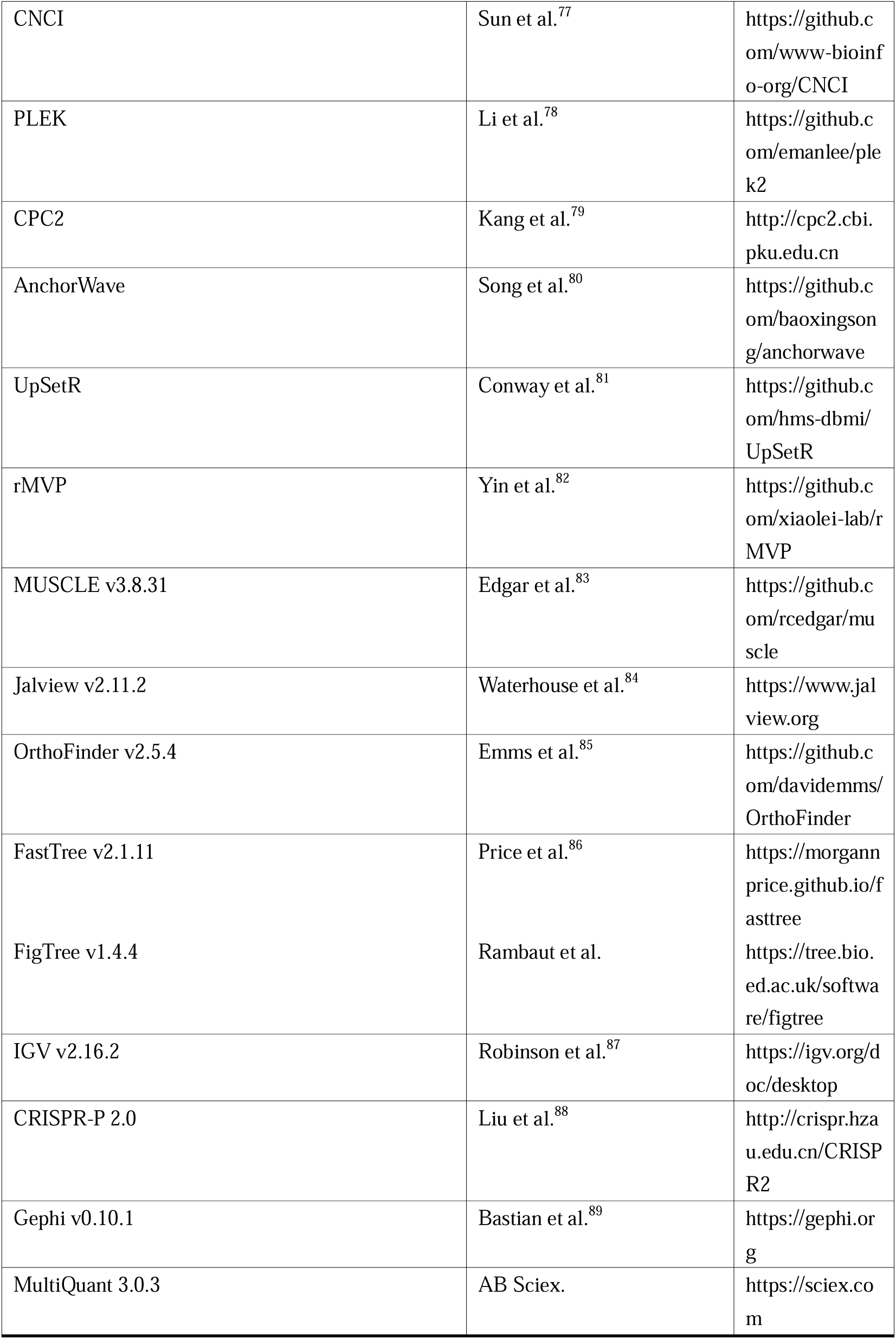

### RESOURCE AVAILABILITY

### EXPERIMENTAL MODEL AND SUBJECT DETAILS

#### Plant materials and growth conditions

CRISPR-Cas9-generated knockout lines of *ZmMp0003340*, *ZmMp0080300*, and *ZmMp0172550* in the KN5585 background, together with their corresponding wild-type controls, were used for phenotypic analysis. Seeds were first germinated in seedling trays, and seedlings at third leaf stage were then transplanted to the field for cultivation with a uniform growth stage. Field trials were performed under standard agricultural practices at the experimental station in Wuhan (30.47°N, 114.35°E) and Sanya (18.15°N, 109.21°E), China.

For etiolation experiments, seeds of the maize inbred line B73 were imbibed in water for 6 h and sown in soil. Plants were initially grown in a controlled environment chamber with a 16 h light (26°C) and 8 h dark (22°C) cycle. After three days, when coleoptiles had reached approximately 1 cm in length, seedlings were shifted to continuous darkness at 25°C for 10 days to induce etiolation before tissue collection.

### METHOD DETAILS

#### Computational pipeline for the identification of microproteins

We established and applied a stringent computational pipeline to identify microproteins across three plant species:

1. Genome-wide sORF prediction: The reference genome of *A. thaliana* (TAIR10), *O. sativa* (IRGSP-1.0), and *Z. mays* (B73_v5) were obtained from Ensembl Plants (release-57) (https://plants.ensembl.org/index.html)^90^. The getorf program from EMBOSS (v6.6.0.0)^56^ was used to predict all potential sORFs encoding 5–100 amino acids, retaining only those with canonical start (ATG) and stop codons.
2. Translatome data processing: Ribo-seq raw data were subjected to quality control with Trim Galore (v0.6.10)^57^, followed by removal of rRNA and tRNA sequences using Bowtie (v2.2.5)^60^. The cleaned reads were subsequently mapped to the respective reference genomes using HISAT (v2.2.1)^58^, with ORF expression profiles quantified by StringTie (v2.1.7)^59^.
3. Proteomics data processing: ORFs with translation expression levels > 1 TPM (from Ribo-seq) were remained for further proteomic analysis. Raw proteomic data were processed using MaxQuant (v1.6.5.0)^63^, retaining peptides with iBAQ > 1 at FDR < 0.01.
4. Species-specific microprotein definition: For *Arabidopsis* and rice, sORFs supported by evidence from both Ribo-seq (> 1 TPM) and proteomic (iBAQ > 1) were defined as high-confidence microproteins.

For maize, step (1) was first used for preliminary screening of potentially translatable ORFs. We further employed a deep learning model (DeepMP, see below) trained on validated microproteins from *A. thaliana*, *O. sativa*, and publicly available peptide sequences to predict potential microproteins in the maize genome. The resulting predictions were then cross-validated against with maize Ribo-seq and proteomic datasets. Only sORFs were both predicted as coding by DeepMp and supported by Ribo-seq and proteomic evidence were designated as high-confidence maize microproteins.

#### Dataset preparation for training DeepMp model

Positive dataset construction: High-confidence, validated microprotein sequences (5–100 aa) from *A. thaliana* and *O. sativa* (as identified above) were combined with experimentally verified peptide sequences obtained from the UniProt and NCBI databases. Peptide sequences were systematically extracted using “peptide” as the search term, applying length restrictions of 5–100 amino acids. All retrieved sequences underwent rigorous manual verification to ensure data quality.

Negative dataset: To construct a high-quality negative set, we selected sequences from the predicted sORFome that exhibited detectable translatome expression but lacked any proteomic support, and showed no significant homology to the positive dataset (BLASTp E-value > 1e^-2^)^64^.

Dataset partitioning: The positive dataset was randomly divided into training (80%) and testing (20%) subsets. To prevent length-based bias, the negative dataset was firstly stratified into six bins according to length (5–15, 15–30, 45–60, 60–75, 75–90, and 90–100 aa) to maintain the natural length distribution across both training (80%) and testing (20%) sets.

#### DeepMp model architecture and hyperparameter optimization

Input sequences were encoded (amino acids 1–20) and standardized to 100 residues via zero-padding^23^. The DeepMp architecture comprised three modules: (1) a 4-layer 1D-CNN block with 3×1 kernels, ReLU activation, and max pooling (kernel size = 2) for local motif detection; (2) a 6-layer Bi-GRU (Bi-directional Gated Recurrent Unit) with 0.2 dropout to model long-range sequence dependencies; (3) a 16-head self-attention mechanism (dropout = 0.2) to weight functionally critical residues^91^. The final classification was performed by a fully connected layer. Key hyperparameters, including the number of layers, learning rate (final: 1e-5), and weight decay (final: 1e-5), were systematically optimized via grid search to maximize predictive performance.

#### Model training and evaluation

The performance of the DeepMp model in microprotein identification was rigorously evaluated using a two-phase training strategy. In the first phase, DeepMp was benchmarked against four representative language models (ALBERT V2^65^, BERT^66^, RoBERTa^67^, and Transformer^91^), sourced from Hugging Face (https://huggingface.co/), using a balanced (positive : negative ≈ 1:1) dataset. Model performance was assessed using accuracy (Acc), sensitivity (Sn), specificity (Sp), and Matthews correlation coefficient (MCC). All models were standardized with identical sequence encoding and output layers, and underwent independent hyperparameter optimization to ensure a fair comparison. In the second phase, the optimized DeepMp model was trained to convergence on an imbalanced (positive:negative ≈ 1:10) dataset to better reflect the expected distribution in genomic data (Figure S1). The final model was applied for genome-wide prediction of putative microproteins in maize.

#### Bioinformatic characterization of Microproteins

Conserved protein domains were identified against the Pfam 32.0^92^ database using hmmscan from HMMER (v3.0)^68^ with an E-value cutoff of 1e^-5^. Signal peptides were predicted with SignalP (v6.0)^69^ (--format none --organism eukarya --bsize 48 --mode fast).

Microprotein loci were classified into five categories (Fig. 2C) based on their positions relative to annotated genomic features (genes, TEs, and lncRNAs) as: intergenic, situated fully in intergenic regions including untranslated regions (UTRs); intronic, fully contained within introns; overlapping, sequences overlapping with any annotated features; contained, entirely within feature boundaries; and antisense, located on the strand opposite to any annotated features. All of the former four classes were located in the sense strand.

Microprotein sequence families were clustered using MMseqs2^70^ with the parameters “--cov-mode 1 -c 0.8 --min-seq-id 0.5”. 3D structures were predicted using AlphaFold 3^93^.

#### Identification of Centromeres

The ChIP-seq data for the CENH3 (NCBI-SRA: SRR21509779) was used to define centromeric regions in maize B73_v5 genome. Clean reads were aligned to the reference genome using Bowtie (v2.2.5)^60^, filtered for high quality (MAPQ > 30) and proper pairing using SAMtools (v1.7)^61^, and PCR duplicates were removed with Picard MarkDuplicates (https://github.com/broadinstitute/picard). Genome-wide chromatin accessibility profiles were then generated by converting the data to bigWig format using bedGraphToBigWig (v1.7)^62^, which enabled the visualization and precise localization of CENH3 enrichment peaks across all chromosomes in the IGV (v2.16.2)^87^.

#### RNA-seq data analysis

Raw RNA-seq reads were processed using fastp (v0.21)^71^ for adapter removal, quality filtering, and base trimming. Clean reads were aligned to the reference genome using HISAT (v2.2.1)^58^ with default parameters. Transcript abundances were quantified as TPM values via StringTie (v2.1.5)^59^ using the reference annotation (GFF3). Tissue specificity was evaluated by calculating the Tau index (range 0–1) using the R package tispec (https://github.com/roonysgalbi/tispec). Co-expression networks were constructed using WGCNA^72^ (power=10, minClusterSize = 100) implemented in R, and visualized in Gephi (v0.10.1)^89^.

#### Genome annotation of TEs, miRNAs, and lncRNAs

Repetitive elements in the maize genome were annotated *de novo* using the EDTA^73^ pipeline (edta.pl) with default parameters. A non-redundant set of maize miRNA was compiled by integrating 465 sequences from PmiREN^74^, 624 from sRNAanno^75^, and 165 from published literature^94,95^, which were subsequently mapped to the reference genome using BLASTn^64^ (-task blastn-short -evalue 1e-2), retaining only the highest-scoring alignments as putative miRNA loci.

lncRNAs were predicted through a stringent filtration workflow. Briefly, transcriptome assembly was performed with StringTie (v2.1.5)^59^ using merged GTF files from 217 RNA-seq samples (Table S3), filtering for length (>200 nt), multi-exonic structure, and class codes (’u’, ‘x’, ‘i’, ‘j’, ‘o’; as assigned by GffCompare^76^). Coding potential was assessed using CNCI^77^, PLEK^78^, and CPC2^79^; only transcripts classified as non-coding by all three tools were retained. Finally, any remaining transcripts with homology to Pfam domains (hmmscan, E-value < 1e^-5^) or the NCBI non-redundant (NR) protein database (BLASTp^64^, E-value < 1e^-5^) were removed.

#### Cross-species conservation analysis

Evolutionary conservation of microproteins and protein-coding genes were evaluated using a two-tiered strategy. First, sequence homology was determined by BLASTp^64^ (E-value < 1e^-5^) against the predicted sORFomes of five Poaceae species (*O. sativa*, *S. viridis*, *S. bicolor*, *Z. mexican*, and *Z. parviglumis*). Second, genomic synteny was performed with AnchorWave^69^ to identify conserved genomic blocks between maize and other species. Microproteins and protein-coding genes exhibiting both significant sequence homology and localization within syntenic regions were defined as conserved. Conservation patterns across species were visualized using UpSetR^81^. The Ka/Ks ratio for each pair was calculated using TBtools with default parameters^96^.

#### eQTL and haplotype analysis

Microprotein expression levels from 368 maize inbred lines (kernel tissues at 15 DAP)^54^ were processed using the RNA-seq data analysis pipeline. Genome coordinates were lifted over from B73 version V4 to V5 using Liftoff^97^. eQTL was mapped with the rMVP^82^ package in R, correcting for kinship and the top five principal components (PCs) estimated by its built-in functions. Haplotypes were defined based on significant eQTL SNPs within the gene body or flanking 2 kb regions.

#### Phylogenetic analysis

Protein sequences were aligned using MUSCLE (v3.8.31)^83^ with default parameters, and the resulting alignments were visualized by using Jalview (v2.11.2)^84^. A species phylogeny was constructed from single-copy orthologs identified across relevant species via OrthoFinder (v2.5.4)^85^. Gene families were aligned with MUSCLE (v3.8.31), and the high-quality alignments were concatenated into a supermatrix. Maximum-likelihood phylogenetic trees were constructed using FastTree (v2.1.11)^86^. The tree was visualized and annotated using FigTree (v1.4.4) (https://tree.bio.ed.ac.uk/software/). Divergence times were estimated from the TimeTree^98^.

#### Validation of microprotein transcription and translation

RNA-seq, Ribo-seq, and proteomic datasets from maize kernels at the grain-filling stage were first integrated to characterize the transcription and translational potential of candidate microproteins. BAM files of the RNA-seq and Ribo-seq were used to confirm transcript coverage and ribosome occupancy at the corresponding loci. We quantified raw proteomic data using MaxQuant (v1.6.5.0)^63^ and assessed peptide support for the candidate microproteins.

The translation of candidate microproteins was further validated using a dual-reporter protoplast assay. Full-length CDS of *ZmMp0003340*, *ZmMp0080300*, and *ZmMp0172550* were cloned in-frame with GFP reporter, under the control of the CaMV 35S promoter. To confirm translation initiation from the microprotein’s start-codon, substitution of start codon mutation (ATG→ATT) was introduced into the microprotein CDS. As a control, a separate mutation was introduced into the downstream GFP start codon (ATG→CTT). Plasmids (10 μg) were transformed into maize mesophyll protoplasts via PEG4000-mediated transformation. Following 16 h incubation at room temperature in WI medium, GFP fluorescence was examined using a Zeiss LSM 880 confocal microscope. Control constructs exhibited robust GFP signals, while experimental mutants completely lost fluorescence, confirming that translation depends on intact start codons.

#### Generation and validation of transgenic lines

Knockout lines were generated via *Agrobacterium*-mediated transformation of maize KN5585 (XINMI Biotechnology Co., Ltd) embryos using the CRISPR-Cas9 system. The sgRNAs targeting candidate genes were designed using CRISPR–P (v2.0) (http://crispr.hzau.edu.cn/CRISPR2/)^88^, and cloned into the pCAMBIA3300 vector under the control of the maize U6 promoter. Gene editing plants were verified by PCR and Sanger sequencing to identify mutant events, and homozygous T2 (for both wild-type and knockout) lines were subsequently selected for phenotypic characterization. All primers used here are listed in Table S4.

#### Quantification of Kernel Amino Acid Content

Kernel amino acid content was determined using LC-MS. For each genotype, mature kernels from 2–3 cobs were pooled and homogenized using a grinding apparatus at 30 Hz for 1 minute. A 100 mg of the resulting powder were transferred to a 2 mL microcentrifuge tube and extracted with 1 mL of methanol (1:5, w/v). The mixture was vortexed thoroughly three times at 10-min intervals, followed by an overnight extraction at 4 °C. Samples were then centrifuged at 12,000 × g for 10 minutes at 4 °C, and the supernatant was filtered through a 0.22 μm syringe filter into amber vials. Metabolomic analysis was performed using an AB Sciex 6500+ QTRAP system coupled with ultra-performance liquid chromatography. Metabolite quantification was conducted with MultiQuant 3.0.3 (https://sciex.com/). Eight biological replicates were analyzed per genotype. The total nitrogen content of 50–110 mg of grain powder was quantified using an Elementar Rapid N Exceed elemental analyzer based on the Dumas combustion method.

### QUANTIFICATION AND STATISTICAL ANALYSIS

Data are presented as mean ± standard deviation (SD). Given that the two groups had equal sample sizes, a two-tailed Student’s t-test was employed to determine statistical significance. For comparisons among multiple groups, one-way ANOVA followed by Fisher’s LSD post hoc test was used, or the Kruskal-Wallis test when appropriate and noted particularly. A P-value < 0.05 was considered statistically significant. All statistical analyses were performed in python. The sample size, number of replicates, and further statistical details are provided in the corresponding figure legends and tables.

