## Supplemental information for "Deep learning reveals a microprotein atlas in maize and uncovers novel regulators of seed amino acid metabolism"

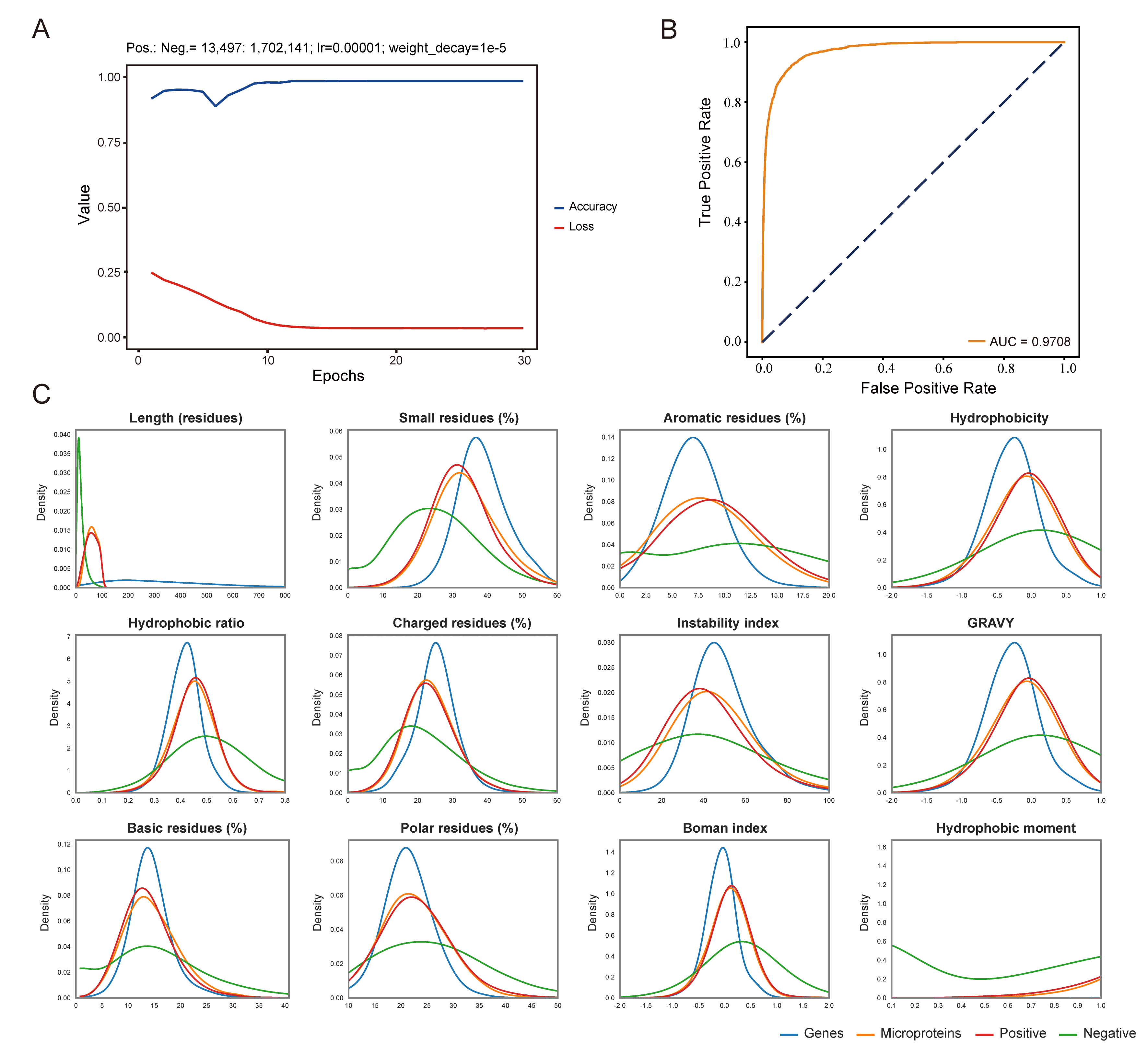

**Figure S1. Performance and validation of the microprotein prediction model.** (**A**) Training curves showing model loss and accuracy per epoch. The model was trained on 13,497 positive and 1,702,141 negative microproteins (using a learning rate of 1e-5 and weight decay of 1e-5). (**B**) The Receiver Operating Characteristic (ROC) curve of the trained model, demonstrating high classification accuracy with an Area Under the Curve (AUC) of 0.9708. (**C**) Density distributions of key physicochemical properties for the gene set, the positive set, the negative set, and the newly predicted microproteins. The predicted microproteins share a highly similar distributions in features (e.g., residue composition, hydrophobicity, and etc.) to the positive set, and both are distinct from the negative set.

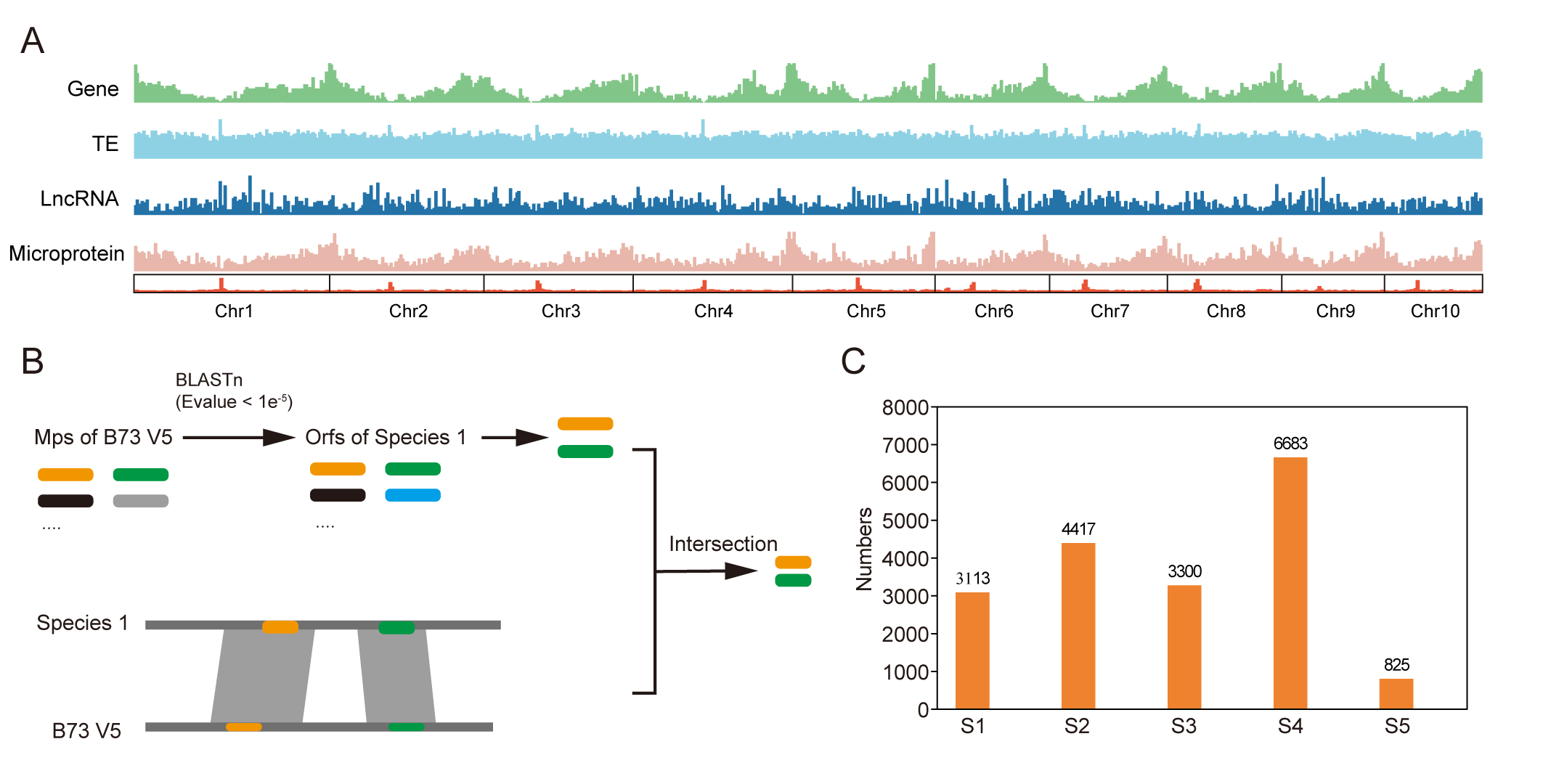

**Figure S2. Genomic distribution and evolutionary conservation of maize microproteins**. (**A**) Chromosomal density of annotated genes, transposable elements (TEs), long non-coding RNAs (lncRNAs), and the predicted microproteins across the 10 maize chromosomes. (**B**) A schematic illustration of the pipeline for identifying conserved microproteins through sequence homology and genomic synteny analysis. (**C**) The distribution of microprotein numbers originating at different evolutionary nodes. Evolutionary strata are defined as: S1 (pre-dating *O. sativa* divergence), S2 (pre-dating *S. viridis* divergence), S3 (pre-dating *S. bicolor* divergence), S4 (*Zea*-genus specific), and S5 (*Z. mays* specific).

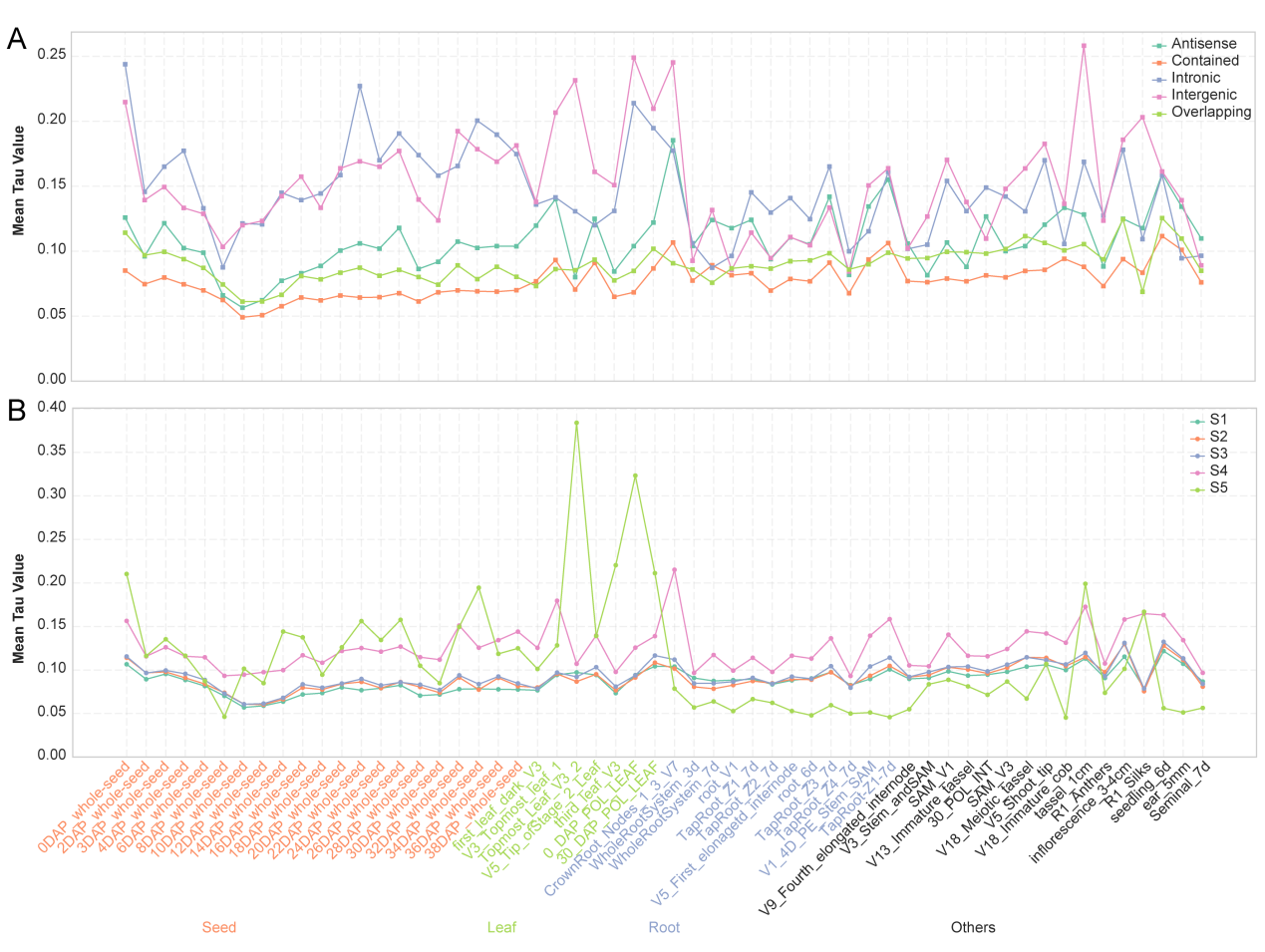

**Figure S3. Tissue specificity of microproteins is associated with genomic origin and evolutionary age.** Tissue specificity (Tau index) was calculated across 56 tissues and developmental stages, which measures the breadth of gene or microprotein expression across analyzed tissues. The Tau index ranges from 0 (housekeeping) to 1 (highly specific). (**A**) Comparison of Tau values for microproteins from different genomic regions, showing that those encoded within intronic and intergenic regions exhibit significantly stronger tissue specificity. (**B**) Comparison of Tau values across evolutionary strata, demonstrating that evolutionarily younger microproteins (S4–S5) are higher tissue specificity.

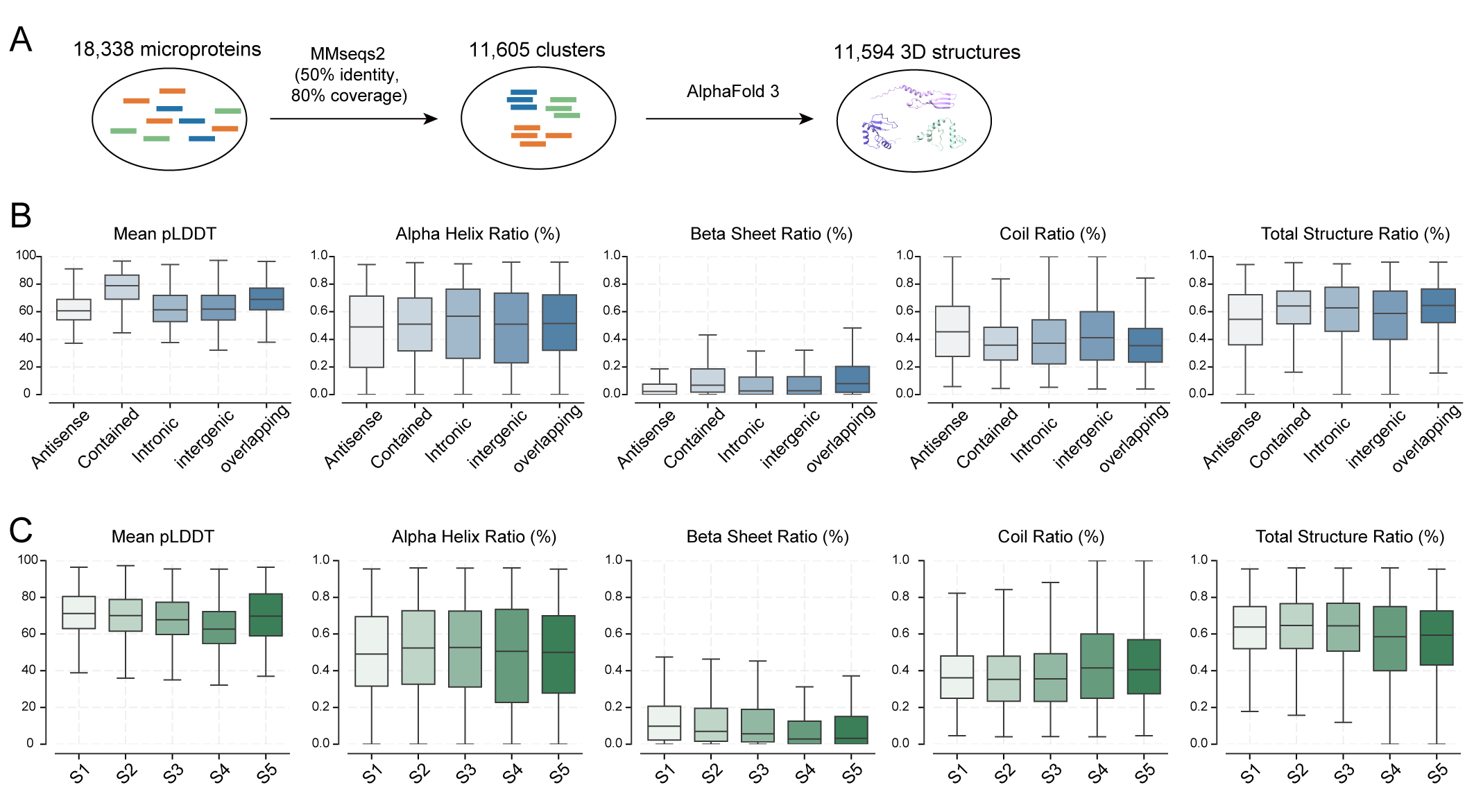

**Figure S4. Structural prediction and properties of maize microproteins**. (**A**) Overview for microprotein clustering and structural prediction. The 18,338 microprotein sequences were clustered, and representative structures were predicted using AlphaFold3. Predictions for 11 microproteins were unsuccessful. (**B–C**) Boxplots comparing structural quality and composition. Metrics include the mean pLDDT (predicted Local Distance Difference Test) score and the ratios of different secondary structures (including Alpha Helix, Beta Sheet, and Coil), as well as the Total Structure Ratio among the predicted structures. Structures are grouped by the genomic regions and evolutionary stratum.
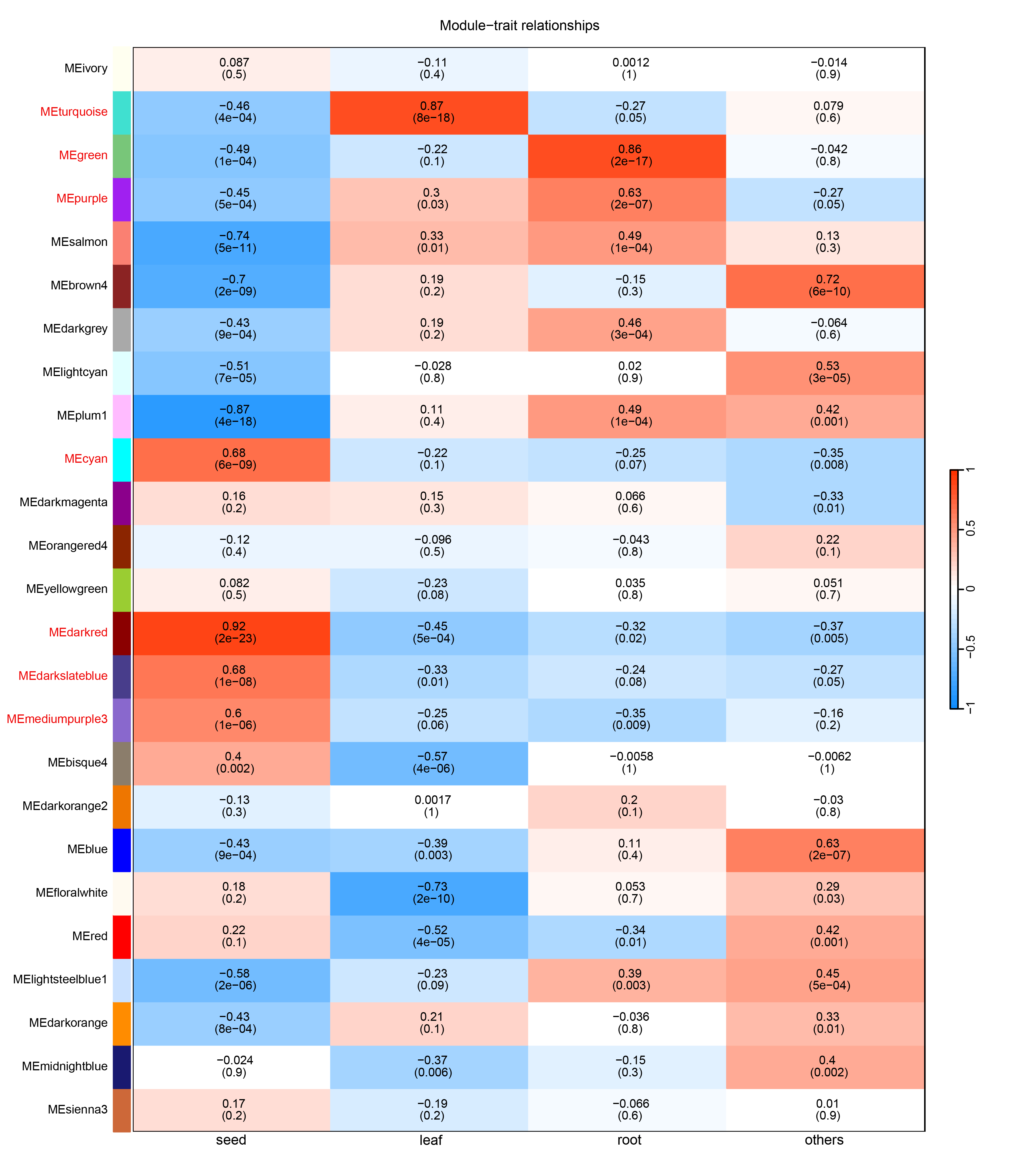

**Figure S5. Module-trait correlation between co-expression modules and maize tissues.** The heatmap depicts the module-trait correlation between gene/microprotein co-expression modules (rows) and specific tissue/developmental stages (columns). Color scale from blue (negative correlation) to red (positive correlation) represents the Pearson correlation coefficient. The corresponding p-value for each correlation is shown in parentheses. Modules highlighted in red were selected for downstream analysis.

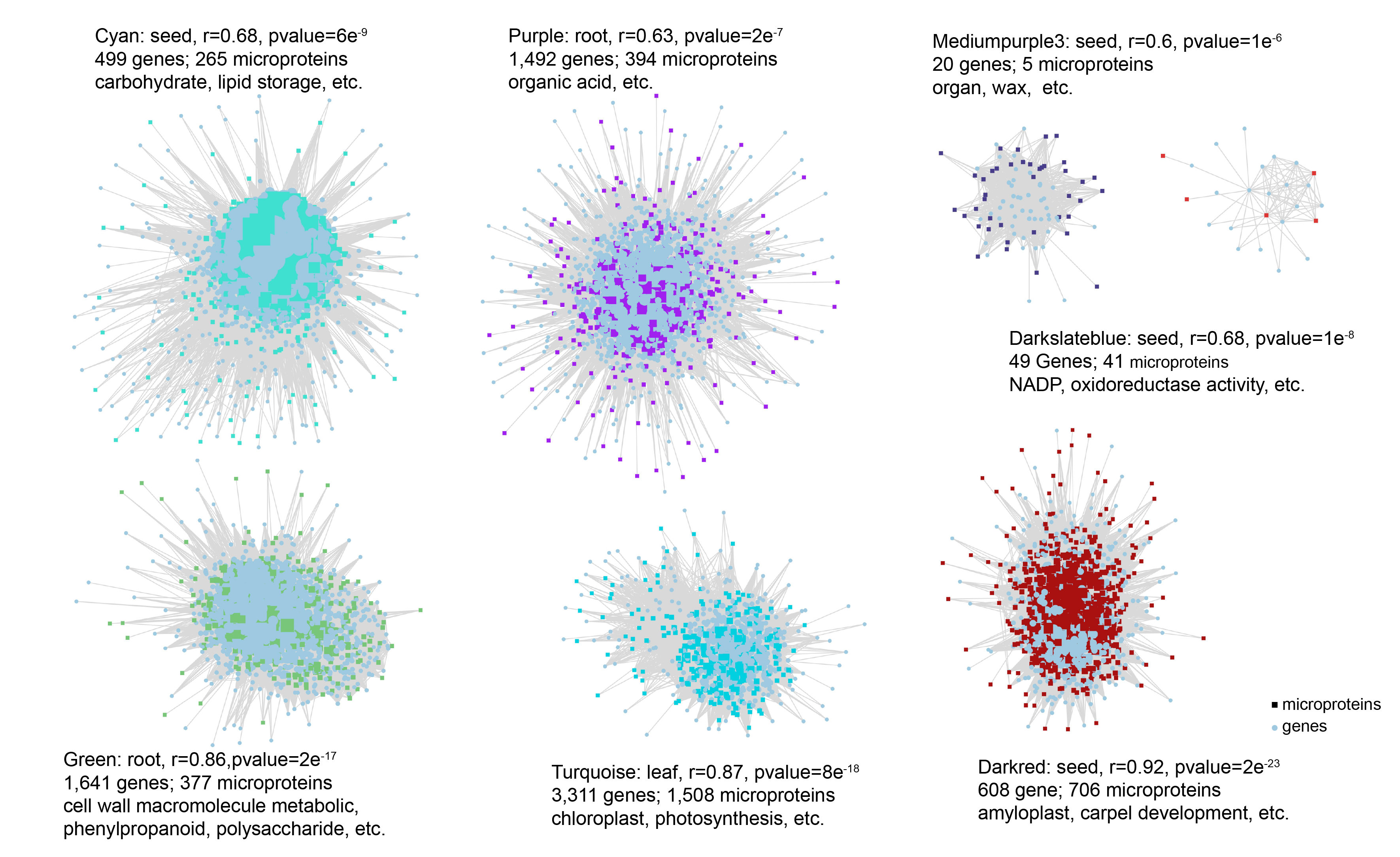

**Figure S6. Co-expression network of microproteins and genes.** Seven co-expression modules significantly associated with the Seed, Leaf, and Root. The Pearson correlation coefficient (r) and associated p-values are indicated adjacent to their respective modules, along with the top enriched Gene Ontology (GO) terms. Genes are represented by light blue circles, and differently colored squares correspond to microprotein modules.

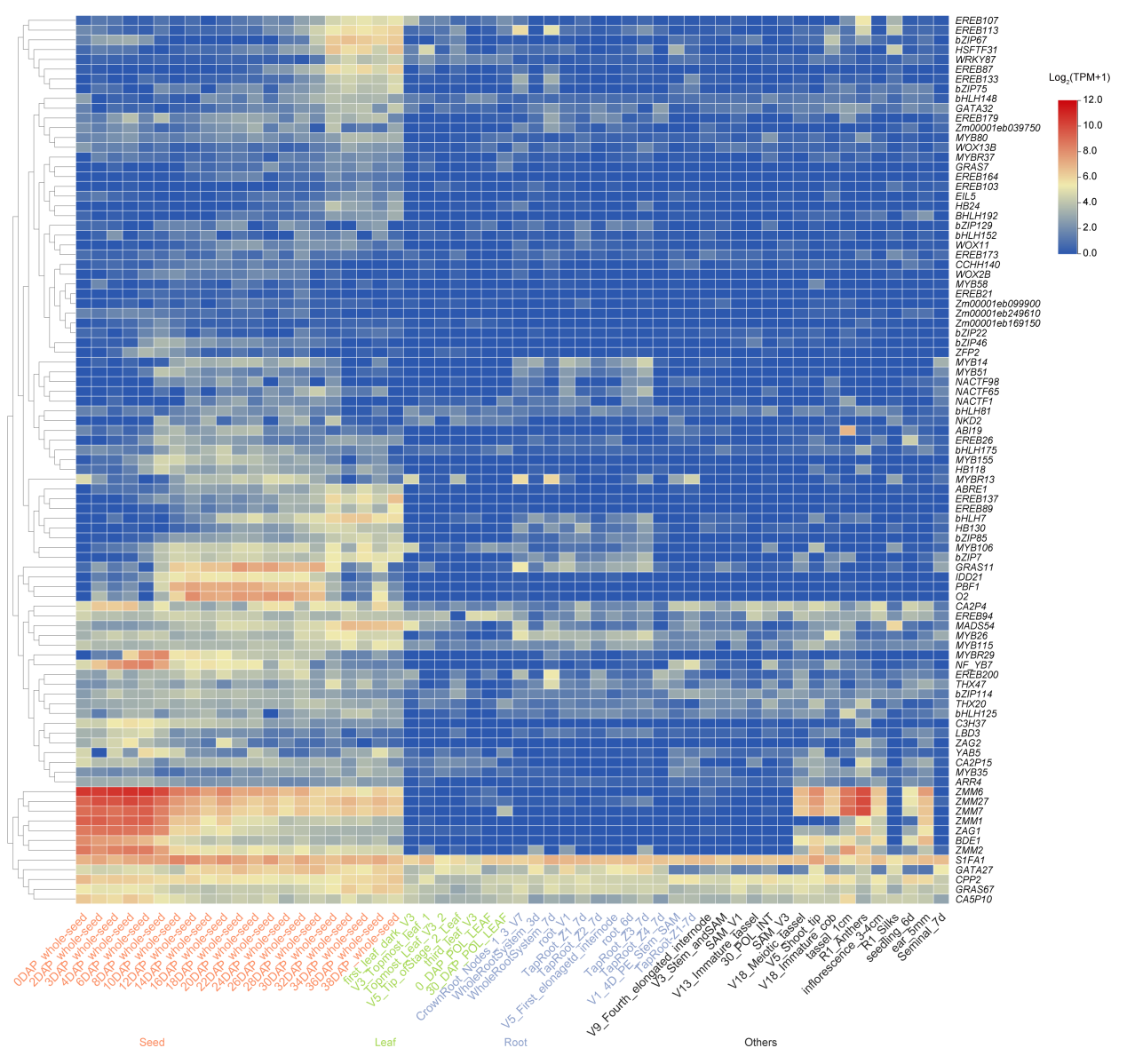

**Figure S7. Expression profiles of transcription factors (TFs) within the seed development-associated co-expression module.** The heatmap displaying log_2_(TPM+1) expression levels of TF genes along the tissue/developmental stages, with red highlights the seed-specific modules.

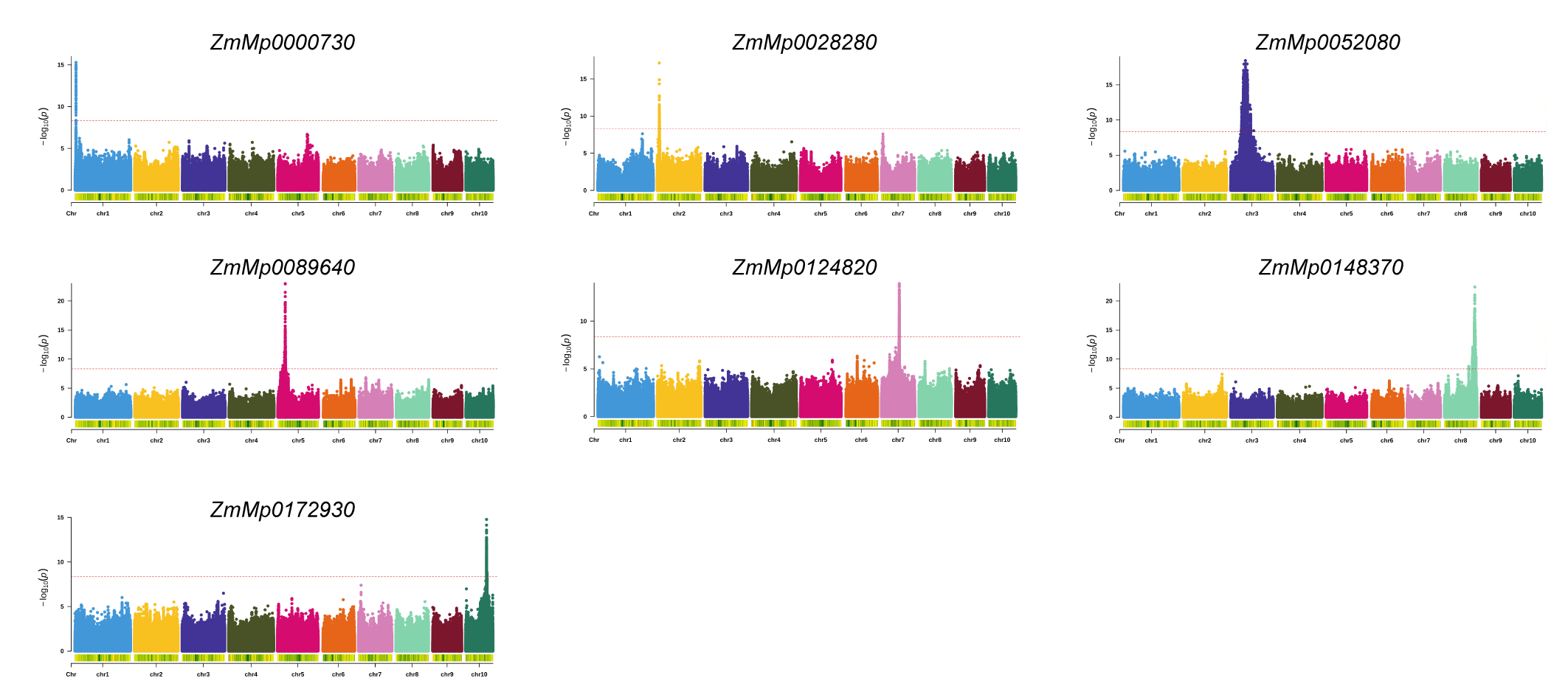

**Figure S8. eQTL analyses for candidate microproteins.** Manhattan plots showing genome-wide associations for the expression levels of seven microproteins. The genome-wide significance threshold (red dashed line) was set by a 5% Bonferroni correction.

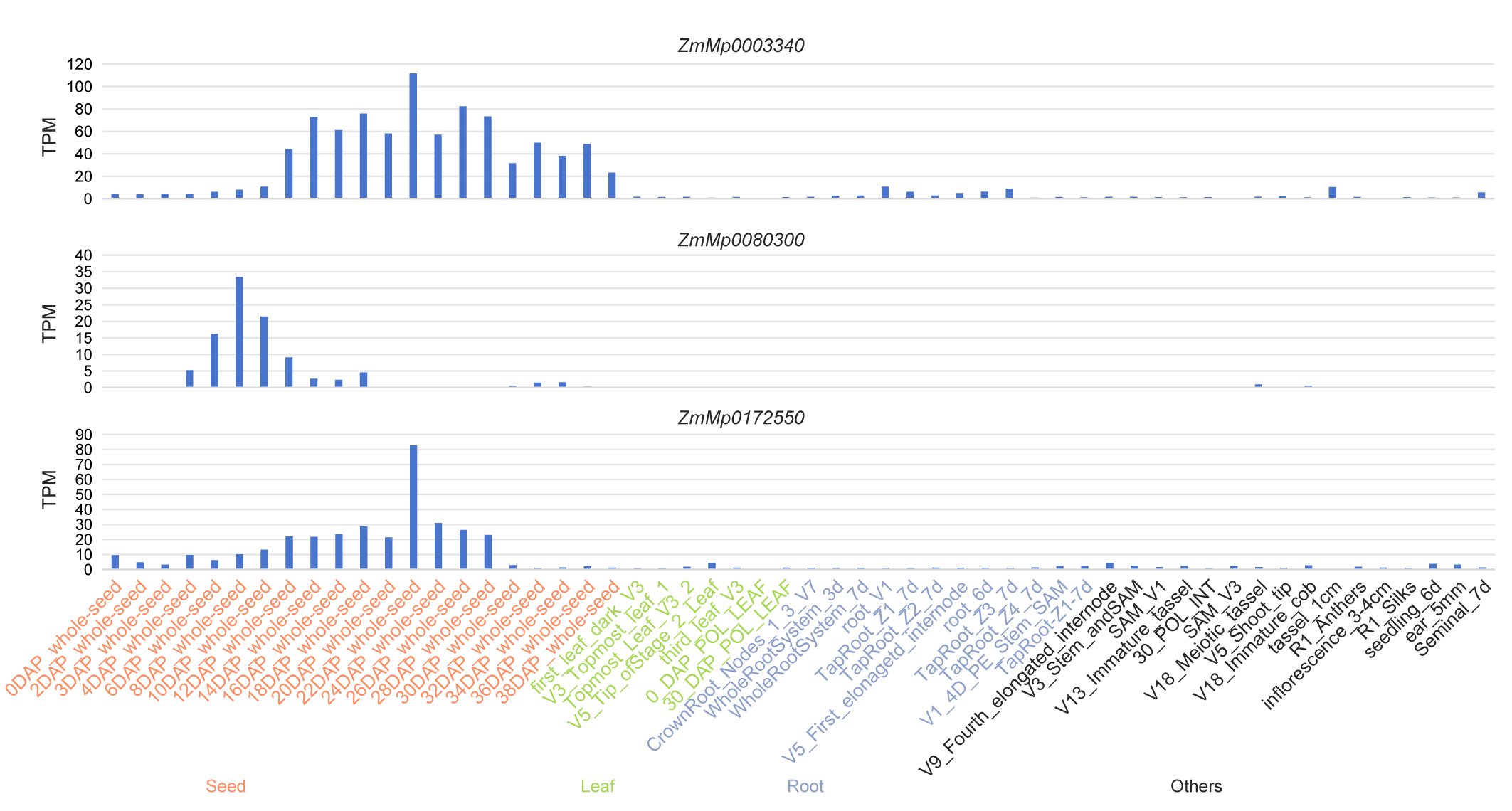

**Figure S9. Transcriptional expression levels of the three microproteins across 56 maize tissues/developmental stages.**

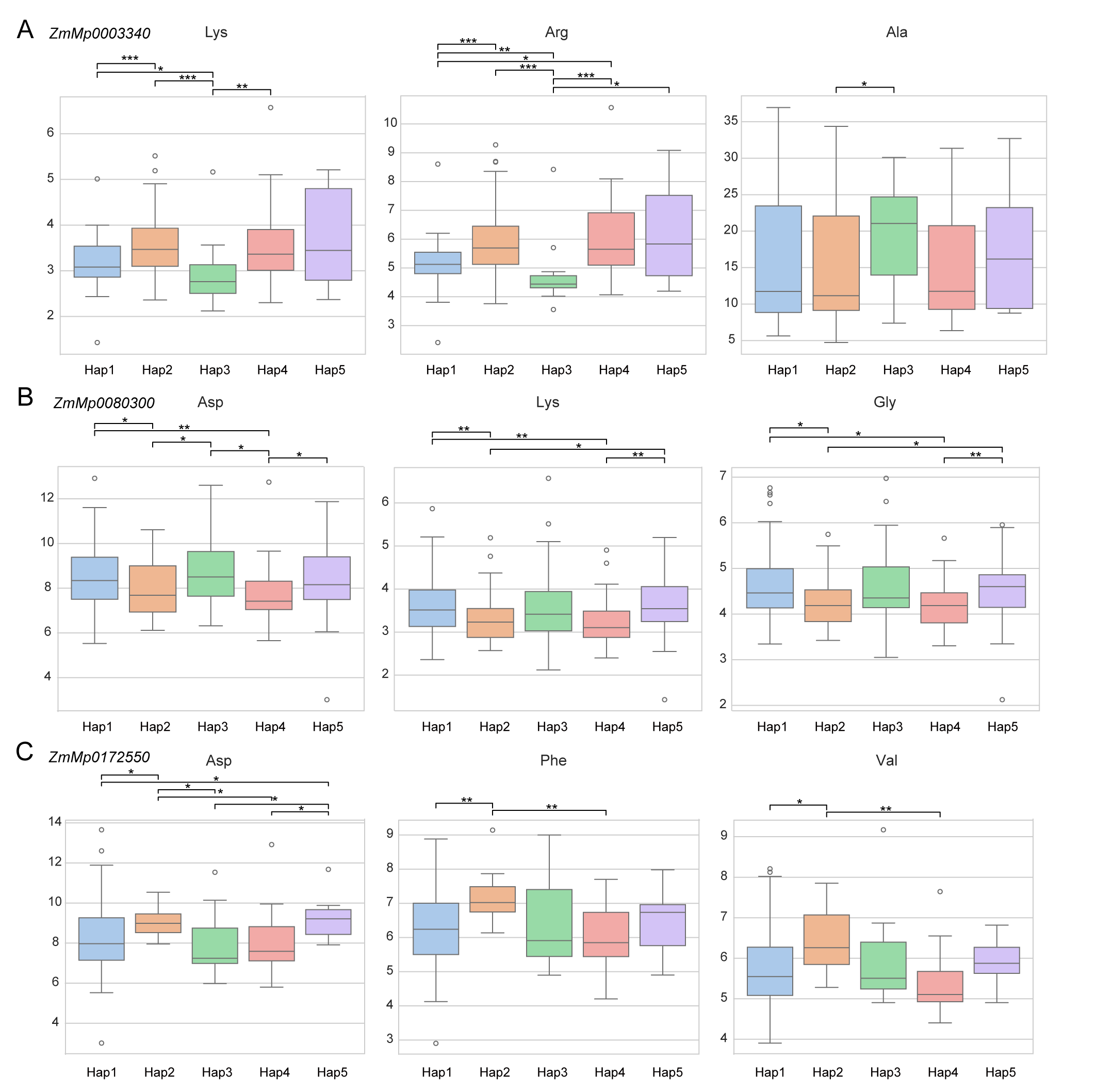

**Figure S10. Haplotype-phenotype association for three microproteins with seed amino acids content.** Association of natural haplotypes of **(A)** *ZmMp0003340*, **(B)** *ZmMp0080300*, and **(C)** *ZmMp0172550* with the relative content of specific amino acids in mature seeds. Data are mean ± SD. Statistical significance was determined by the Kruskal-Wallis test (*P < 0.05, **P < 0.01, ***P < 0.001). The sample size for each haplotype is detailed in Figure 4E.

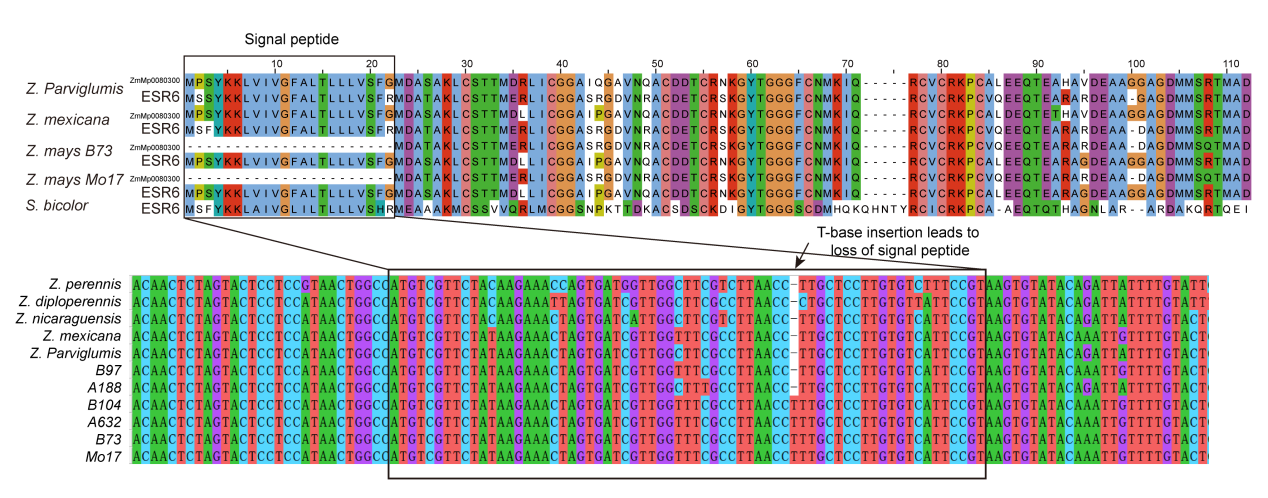

**Figure S11. Multiple sequence alignment of ZmMp0080300 and its homolog ESR6. (Top)** The amino acid alignment of ZmMp0080300 and its previously identified homolog Embryo Surrounding Region 6 (ESR6), and (**Bottom**) the corresponding genomic DNA alignment of the signal peptide region from 11 *Zea* accessions. A single T-base insertion is responsible for the loss of the signal peptide (that causing a frameshift) in certain maize inbreds.

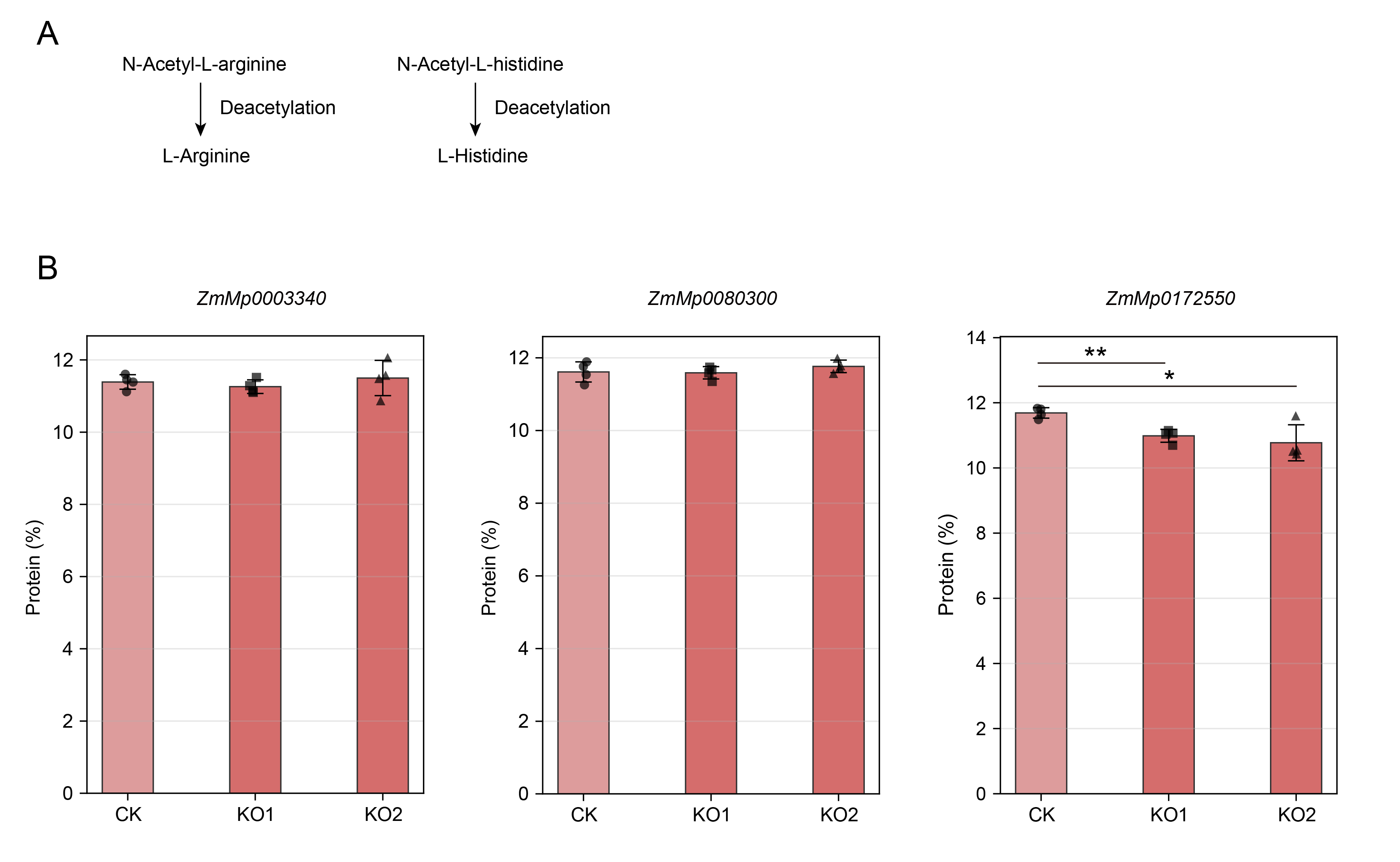

**Figure S12. Conversion of N-acetylated intermediates to amino acids and functional validation of *ZmMp0172550* affecting seed protein content. (A) Schematic showing the conversion of N-acetyl-L-arginine and N-acetyl-L-histidine into L-arginine and L-histidine, respectively, through deacetylation reactions. In the arginine pathway, due to the limited sensitivity of LC-MS for L-arginine detection, its precursor N-acetyl-L-arginine was measured instead, allowing indirect assessment of potential changes in L-arginine levels. N-acetyl-L-arginine can be converted into L-arginine through deacetylation. In contrast, both N-acetyl-L-histidine and L-histidine were directly detectable by LC-MS in the histidine pathway. (B) Measurement of total protein content in** mature seeds of **wild-type (WT) and knockout (KO) of the three** microproteins in the **KN5585 background. The knockout of *ZmMp0172550* resulted in a significant decrease in total protein content. Data are mean ± SD (n=4)**. Significance was determined by a two-tailed Student's t-test (*P < 0.05, **P < 0.01, ***P < 0.001).

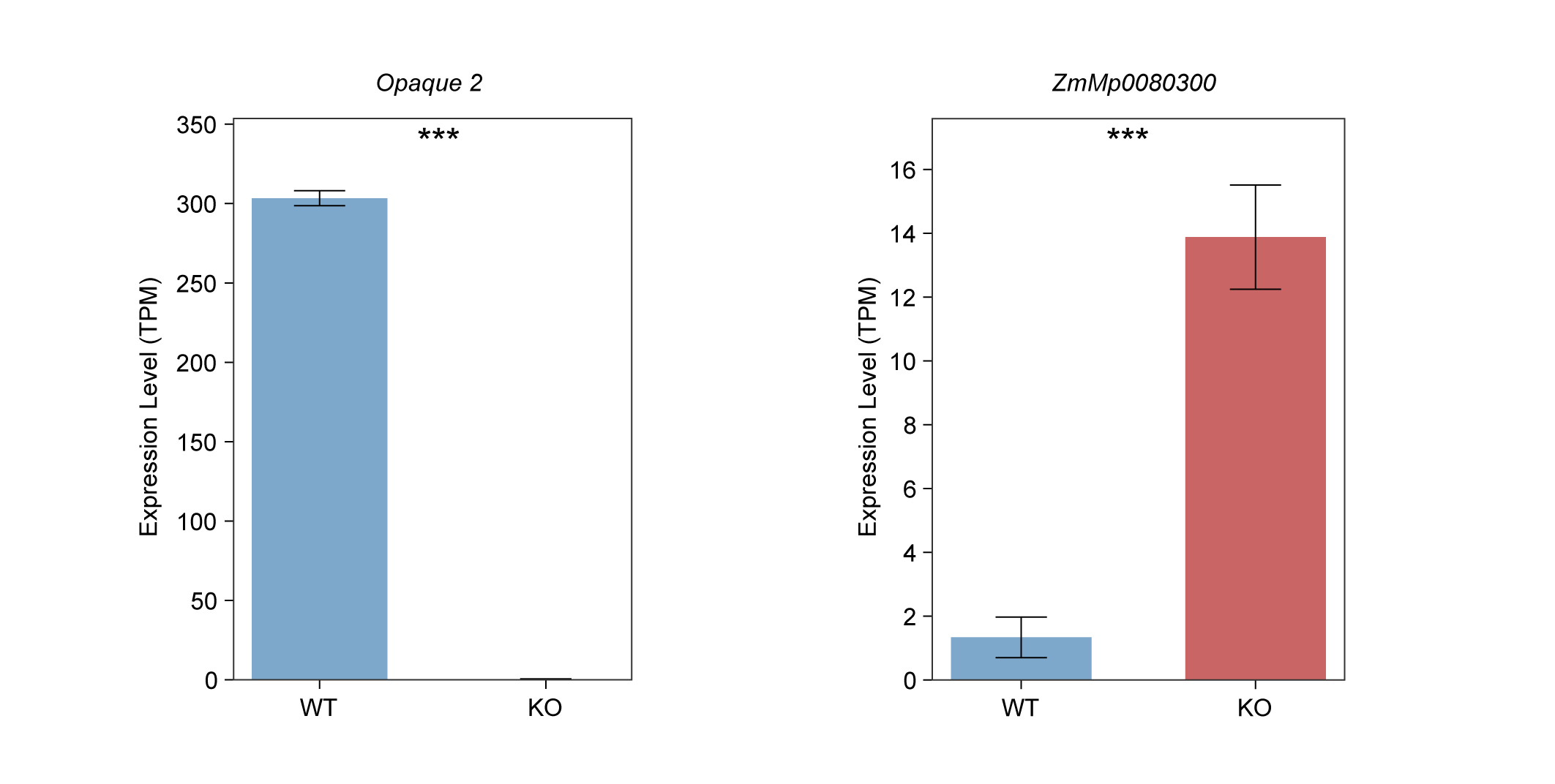
**Figure S13. Regulatory relationship between *Opaque2* and *ZmMp0080300*. Expression values (TPM) of *Opaque2* (left panel) and *ZmMp0080300* (right panel) in** wild-type (WT) and ***Opaque2*-KO background.** The loss of *Opaque2* expression in the KO line confirms the knockout. The concomitant change in *ZmMp0080300* expression suggests it is regulated by, or acts downstream of, the Opaque2 transcription factor. Data are mean ± SD (n=3). Significant differences were determined by a two-tailed Student's t-test (*P < 0.05, **P < 0.01, ***P < 0.001).
